# Nuances of floral mimicry in Queen of Sheba orchids

**DOI:** 10.1101/2025.03.13.642682

**Authors:** Daniela Scaccabarozzi, Gavin R. Flematti, Mark Brundrett, Mario Vallejo-Marín, Klaus Lunau, Monica Gagliano, Emiliano Pioltelli, Terry Houston, Andrea Aromatisi, Nina Sletvold

## Abstract

- **Background and Aims** Floral deception can range from finely tuned mimicry of specific rewarding plants (Batesian mimicry) to a general resemblance of the rewarding community (generalised food deception). We focused on two species in the deceptive *Thelymitra variegata* complex (Queen of Sheba orchids) to examine whether pollination is consistent with generalised food deception or with specific mimicry of co-flowering buzz-pollinated tinsel lilies (*Calectasia* spp.).
- **Methods** We assessed i) overlap in flower colour and scent between orchids, tinsel lilies, and other flowers in the rewarding community, ii) pollinator sharing between orchids, tinsel lilies and other species, and ii) quantified orchid reproductive success in relation to abundance of tinsel lilies and other rewarding species.
- **Key Results** We found that *T. speciosa* matches flower colour and scent of its co-flowering tinsel lily, *C. gracilis*, while *T. porphyrosticta* partly overlaps with co-flowering *C. demarzii* in both traits. The orchids and tinsel lilies shared flower visitors, and the only visitor observed to transfer orchid pollen was *Anthoglossa plumata* bee (Colletidae) that displayed a behaviour associated with buzz pollination. This species was also observed to buzz-pollinate tinsel lilies. In *T. speciosa*, fruit production increased with abundance of both tinsel lilies and other rewarding species with similar flower colour, suggesting a combination of mimicry and magnet effects. However, relationships were driven by a single population. There was no association between rewarding community and reproductive success in *T. porphyrosticta*.
- **Conclusions** The results are partly consistent with mimicry towards the co-flowering tinsel lily for both orchids. Specific mimicry is strongly supported by a close resemblance involving multiple floral traits, and by bee behaviour associated with the buzz-pollinated model. However, this contrasts with the lack of consistent effects of tinsel lily abundance on orchid reproductive success. The findings further suggest that shared floral scent may reinforce visual similarity and highlight that scent may be important in food mimicry systems. More generally, the *T. variegata* complex offers an interesting system for studying nuances between specific mimicry and generalised deception.

## INTRODUCTION

Most flowering plants produce nectar or consumable pollen to reward their pollinators, but several species in at least 32 plant families (Renner, 2006) lack any form of reward and are classified as deceptive. Floral deception is a particularly common reproductive strategy in the Orchidaceae, with around 40% of species globally recognised as deceptive (Renner, 2006; Ackerman *et al*., 2023). These species do not produce any nectar, and their pollen is packaged into pollinia, which cannot be used as a food source by pollinators. Most deceptive orchids are food deceptive and attract pollinators by mimicking the floral traits of co-flowering species that do offer rewards. These species can exploit the food-seeking behaviour of pollinators through a general resemblance to rewarding species, known as generalized food deception (Dafni, 1983), or imitate the flowers of a specific model species that provides a reward, which is known as Batesian food-source mimicry (Jersáková *et al*. 2009; Johnson and Schiestl, 2016). In between these two strategies, some food-deceptive plants also resemble an entire guild of model species (Dafni, 1983; Bernhardt and Burns, 1986; Dafni and Calder, 1987; Indsto *et al*., 2006; Scaccabarozzi *et al*., 2018; 2020). This suggests a continuum from generalized food deception to Batesian mimicry (Jersáková *et al*., 2009; Johnson and Schiestl, 2016), where the two extremes differ in the existence of a specific model species and the degree of pollination specificity.

In cases of Batesian mimicry, model and mimic species are expected to occupy the same habitat, exhibit overlapping flowering periods, and share flower signals and pollinators, and the pollinator(s) should perform a behaviour associated with its model plant (Roy and Widmer, 1999; Johnson and Schiestl, 2016). In addition, the presence of the model should increase the mimic’s fitness (Dafni, 1983). Many studies have suggested mimicry based on overlapping distribution and phenology but less is known about the specific traits that convey similarity, and whether orchid fitness is affected by the presence of putative model species (Jersáková *et al*., 2009). In general, convergence on multiple flower signals could be expected to maximise the probability of pollinators confusing models and mimics. For example, the South African orchid *Disa pulchra* closely resembles *Watsonia lepida* in terms of flower colour, inflorescence architecture, and nectar guides (Jersáková *et al*., 2012), suggesting that a close visual match is necessary for successful mimicry. In contrast, *D. pulchra* does not share scent profile with the model species (Jersáková *et al*., 2012). Similarly, *Orchis israelitica* was found to resemble its putative model, *Bellevalia flexuosa,* in terms of colour, but not scent (Galizia *et al*., 2005). These results indicate that visual convergence may be sufficient in food-deceptive systems, although they contrast with a bee behavioural experiment suggesting that specialised mimics would benefit from sharing scent with its model (Gumbert and Kunze, 2001). It is clear that more studies that focus on both visual and olfactory signals are needed to determine the relative role of these signals in mimicry. Only a few studies have documented that increasing abundance or proximity of the model species benefits mimic fitness (Anderson and Johnson, 2006; Peter and Johnson, 2009; Duffy and Johnson, 2017), indicating adaptive resemblance.

In cases of generalised food deception, deceptive species are expected to overlap in flowering period and share pollinators with the rewarding species in the community, similar to Batesian mimicry. However, unlike Batesian mimicry, generalised species do not benefit from resemblance to specific model plants, but instead exploit common floral signals shared by multiple co-flowering plants and rely on the generalist foraging behaviour of their pollinators (Cozzolino *et al*., 2005; Jersáková *et al*., 2012; Peter and Johnson, 2013; de Jager and Anderson, 2019). In other words, generalised deceptive orchids benefit from a ‘general search image’ of food-seeking pollinators (Goulson, 2000; Johnson and Schiestl, 2016), and their colour, scent and morphology is expected to be broadly appealing rather than closely matching any particular rewarding species (Jersáková *et al*., 2006). Consequently, fitness of a generalised species is not expected to increase with presence or abundance of any specific rewarding species in the community, but rather with the overall abundance of rewarding plants (magnet species effect; Thompson, 1978; Johnson et al., 2004; Jersáková et al., 2016). Still, some studies indicate that the probability that a pollinator will visit a generalised deceptive plant increases with flower colour similarity to co-flowering rewarding species (Gumbert and Kunze 2001; Johnson et al. 2004), suggesting that a nonspecific colour resemblance could be beneficial. With a likely continuum from generalized food deception to Batesian mimicry, no single criterion should be used to define the deceptive strategy. Instead, inferences on strategy will need to combine information on the precision of trait matching and pollinator behaviour, and on the relationship between fitness of the deceptive species and abundance of the overall rewarding community versus abundance of putative model species.

The Southwest Australian Floristic Region (SWAFR) is a biodiversity hotspot that hosts several orchid species in the genus *Thelymitra,* commonly known as sun orchids. Within *Thelymitra*, a group called Queen of Sheba orchids (*T. variegata* species complex) are believed to exhibit food mimicry, superficially resembling pollen-rewarding tinsel lilies (*Calectasia* spp.) that co-flower with the orchids (Adams and Lawson, 1993; Jeanes, 2009). However, it remains unknown whether pollination is facilitated by resembling these specific model plants, i.e., Batesian mimicry (Jersáková *et al*., 2006), or by resembling rewarding plant species in general, i.e., generalized food deception (Jersáková *et al*., 2012; Johnson and Schiestl, 2016). Here, we specifically examined whether pollination of the *T. variegata* group is more consistent with generalized food deception or with specific mimicry of tinsel lilies (*Calectasia* spp.), focusing on two relatively common species in the group. The two orchid species are similar in terms of flower colour and morphology but differ in putative model species. If mimicry is specifically addressed towards tinsel lilies, we expect that *Thelymitra* and *Calectasia* i) share visual (morphology and colour) floral traits that are not shared with other rewarding species; ii) share scent profiles that are not shared with the remainder of the flowering community; and iii) have specific pollinators that display the same behaviour on the orchids and tinsel lilies. In addition, we expect iv) orchid reproductive success to increase with abundance of tinsel lilies. If generalized food deception is involved, we expect that i) orchid visual floral traits are common among co-flowering plants but not specific to any one species; ii) scent is a general attractant and does not resemble a particular co-flowering species; iii) orchids attract pollinators that display a general foraging behaviour not specifically induced by any particular species; v) orchid reproductive success increases with overall abundance of the co-flowering rewarding community, or with abundance of rewarding species with similar flower colour.

## MATERIALS AND METHODS

### Study species

*Thelymitra* is a primarily Australian genus of terrestrial geophytes in the Orchidaceae, comprising approximately 130 recognised species. Most species are considered to be food-deceptive, but the genus includes some autogamous species (Brundrett, 2024). Plants produce a solitary scape in any given year (Jeanes, 2009), and flowers open on sunny days with temperatures above 18°C (Brown, 2022). Within the *T. variegata* complex, five taxa share a similar flower morphology, and are all believed to mimic *Calectasia* flowers in shape, colour, and reproductive structures (anthers), to achieve pollination by pollen-consuming native bees (Bower, 2001; Jeanes, 2009; Brown, 2022). Populations are generally few, small, and widely dispersed. The taxonomic status of the group is currently unresolved, and whether taxa should be recognised as ecotypes or distinct species remains to be clarified (Jeanes, 2009).

We focused on two taxa, *T. porphyrosticta* and *T. speciosa,* hereafter referred to as species, with adjoining distribution and overlap in flowering phenology and habitats. Both species co-flower with a similar assemblage of rewarding plants and are likely to hybridize where distributions overlap (Edens-Meier *et al*., 2014; Brown, 2022). The Southern Queen of Sheba, *T. porphyrosticta*, grows in sandy soils in damp, open shrublands and woodlands (Sandiford and Barrett 2010), and flowers with up to five purple and violet flowers during August and September (Jeanes, 2009; Brown, 2022). Due to its rarity and narrow distribution, the species is listed as priority flora in Western Australia (Florabase, 1998). The Eastern Queen of Sheba, *T. speciosa*, is a more widespread species that grows in sandy-clay soils in shrublands and exposed plains, and flowers with one or more purple and orange flowers from late June to September (Jeanes, 2009; Brown, 2022). The species is not considered threatened in Western Australia, but populations are often very small (< 10 individuals) and isolated (separated by 50-200 km) (Florabase, 1998). Both orchid species have a star-shaped flower with a golden-yellow column with prominent pseudo-anther appendages that protrude conspicuously from the centre of the flower (Fig. 1). Pseudo-anthers typically resemble anthers in flowers pollinated by pollen-feeding insects (Dafni and Bernhardt, 1989; Lunau *et al*., 2021), and column appendages of Queen of Sheba orchids closely resemble anthers of *Calectasia* species (Fig. 1). Both orchid species are self-compatible but require an external vector for pollination (Jeanes, 2009). Fruit production within a given flowering season is strongly pollen-limited, with most or all flowers on a scape forming fruit after pollination by hand (Brundrett, 2019).

**Fig. 1.**
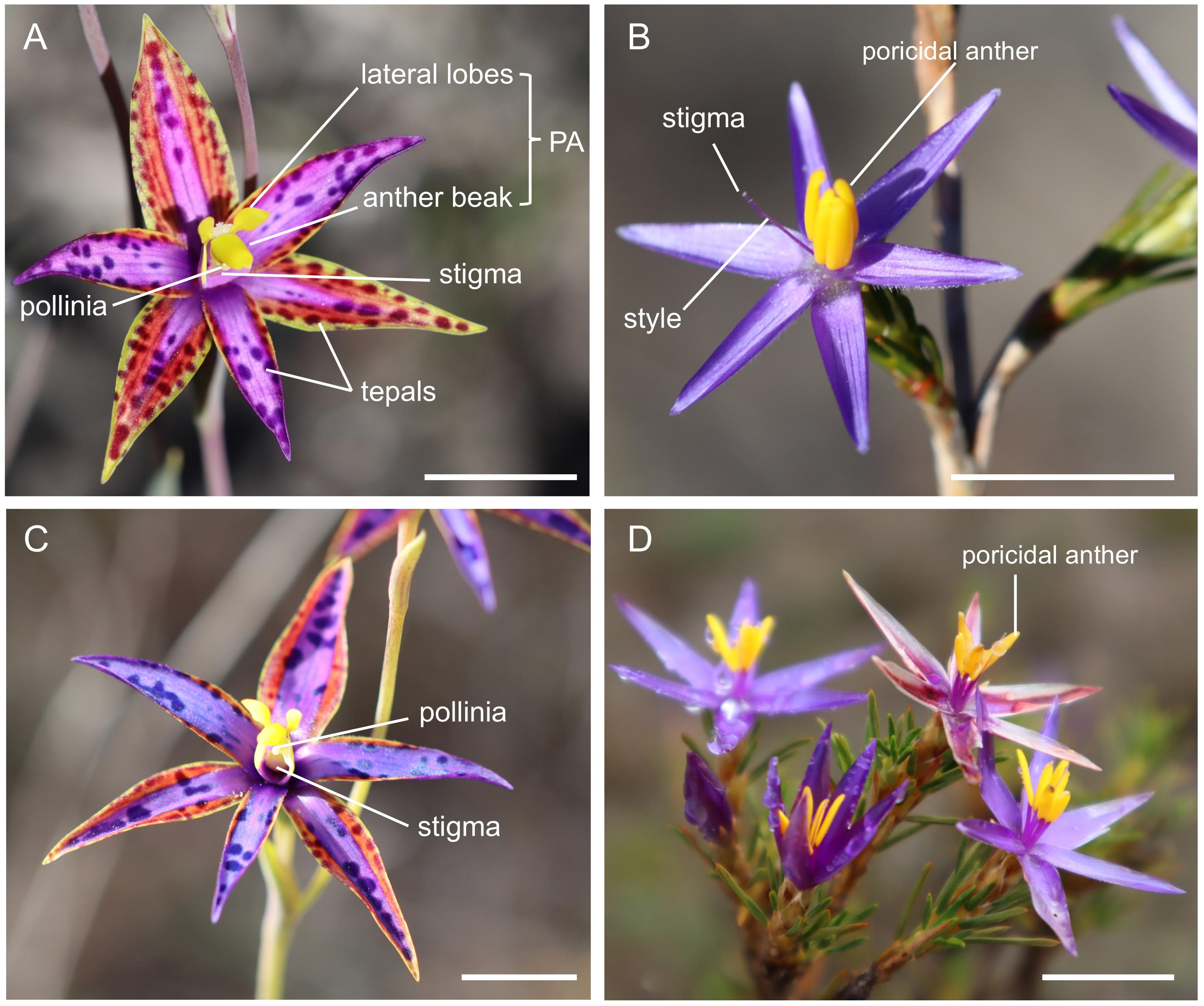
Flower of **A:** *Thelymitra speciosa* (Orchidaceae), indicating the placement of the stigma, the two pollinia attached to a single viscidium, and the pseudoanther (PA: pseudoanther or false anther, consisting of the lateral lobes and anther beak; Jeanes, 2009); **B**: *Calectasia gracilis*; **C**: *T. porphyrosticta,* and **D**: *C. demarzii*. Scale bar: 1 cm. Photo credits: Daniela Scaccabarozzi

### Study sites

We collected data on *T. speciosa* from two populations in 2020 and from four populations in 2021-2022, all situated within the Tozer’s Bush Camp, in proximity to Fitzgerald River National Park. We collected data on *T. porphyrostricta* from two populations in 2021 and from five in 2022, one in Waychinicup National Park in both years, and from one (in 2021) to four (in 2022) in Gull Rock National Park. A habitat description for all populations is presented in [**Supplementary information**, Table S1]. Both orchids often co-occur and co-flower with putative model species in the genus *Calectasia*, tinsel lilies. Tinsel lilies are small, erect shrubs, with a multitude of small, blue to purple, star-shaped flowers, with conspicuous yellow anthers. In the Tozer’s Bush Camp, *T. speciosa* co-flowers with *Calectasia gracilis* and, in Waychinicup National Park, *T. porphyrosticta* occurs with *C. demarzii* (Fig. 1). In Gull Rock National Park, *C. demarzii* has previously been recorded (Florabase, 1998), but there were no flowering tinsel lilies in the focal orchid populations during the study years. Across all study sites, *Calectasia* individuals were sporadic, with a patchy distribution.

Both orchids also co-flower with several other rewarding species, that could provide a magnet effect [see **Supplementary Information**, Table S1]. This includes species that to the human eye share flower colour with the orchids (*Andersonia caerulea, A. simplex, Conospermum caeruleum, Dampiera linearis, Melaleuca suberosa*), as well as several other species [**Supplementary Information**, Table S1]. Plants were identified using Florabase (1998), and by consulting experts (Russell Barrett at Botanic Gardens of Sydney; Sarah Barrett at Department of Parks and Wildlife of Western Australia; Paul Armstrong at Wildflower Society of Western Australia).

### Floral trait similarity between orchids, tinsel lilies, and other rewarding species

To characterize floral trait similarity between the two orchids (*T. speciosa*, *T. porphyrostricta*), the two putative model species, the tinsel lilies (*C. gracilis, C.demarzii*), and other co-flowering rewarding species, we measured flower morphology, colour and scent in the Tozer’s Bush Camp for *T. speciosa* and its associated rewarding species, and in Gull Rock National Park for *T. porphyrosticta* and its associated co-flowering plants [**Supplementary information,** Table S1]. We measured all traits on the orchids and tinsel lilies. For the remaining community, we limited the colour comparison to the most common co-flowering species that to the human eye were similar in colour to *Thelymitra* spp. (*A. caerulea*, *A. simplex*, *C. caeruleum, D. linearis, M. suberosa)* [**Supplementary Information,** Fig. S1], but also included two common species with contrasting colour, i.e., yellow-orange *Daviesia hakeoides,* and yellow *Petrophile squamata*. We limited the scent comparison to the most common co-flowering species with perceived scent (*A. caerulea*, *A. simplex, D. hakeoides, Hibbertia psilocarpa, Leptospermum laevigatum*, *Leucopogon glabellus*, *M. suberosa*) [**Supplementary Information,** Fig. S1]. Except for *Calectasia*, none of the included co-flowering plants had poricidal anthers and were buzz pollinated.

#### Morphology

For both orchid and tinsel lily species, we measured traits predicted to influence pollinator attraction (plant height, corolla width, corolla height, corolla symmetry, presence of exposed pollen, stamen/pseudoanthery height) on 10 individuals. We also obtained trait values from the literature (Marchant *et al*. 1987) for 16 other co-flowering species that occurred at the study sites [**Supplementary Information**, Table S2]. We further classified co-flowering species based on reward type (only pollen, both pollen and nectar, only nectar) according to Marchant *et al*. (1987) and Brundrett *et al*. 2024 [**Supplementary Information**, Table S2]. Morphological floral trait similarity between *Thelymitra* spp*., Calectasia* spp, and other co- flowering plants was visualised by Non-Metric Multidimensional Scaling analysis (NMDS) using the function *meta*MDS in the vegan package in R (Oksanen *et al*., 2013).

#### Colour

To characterise flower colour similarity, we measured spectral reflectance on flowers (tepals or sepals) from six individuals of each included species, randomly collected at the field sites. Measurements were taken on the whole surface of the petals as diffuse reflectance. We used a spectrophotometer (Cary 4000 UV–VIS spectrophotometer; Agilent Technologies) with integration time of 50 ms, and a tungsten halogen visible light source and a deuterium arc UV light source. Measurements were taken across the entire tepals or sepals surface as diffuse reflectance. To estimate how pollinators perceive colours, we used the conventional bee colour vision model (Chittka, 1992; Chittka and Kevan, 2005) which is based on the sensitivities of photoreceptors of the honeybee, *Apis mellifera* (Chittka, 1992). To quantify colour similarity between species, we calculated the mean Euclidean distance between colour loci coordinates in the hexagon (n=6 per species). In addition, we applied false colour photography in bee view as per Verhoeven *et al*. (2018) and Lunau *et al*. (2021) to identify potential specific colour patterns shared by orchids and tinsel lilies.

#### Scent

To assess scent similarity, we analysed the scent from a single flower from each of five individuals of the two orchids and from each of four individuals of the seven co-flowering species with perceived scent (*A. caerulea*, *A. simplex, D. hakeoides, H. psilocarpa, L. laevigatum*, *L. glabellus*, *M. suberosa*). Each flower was placed in a headspace vial (Supelco, 20 mL, SU860097 with magnetic screw septum caps, p/n SU860101) and immediately stored in a portable car freezer at -18°C. The sampling was conducted during the period of peak activity of potential pollinators, i.e., when floral scent production is expected to be at its maximum. To control for damage-induced scent emission we picked sections of stems together with each flower (pedicel included) sample and placed the stem parts in separate vials for subsequent analysis. Four air samples from each site were taken as controls. Peaks present in control samples, or both found in stem and flower samples were not considered part of the floral scent profiles.

Within one week after sampling, we analysed scent samples of flowers and stems (samples were defrosted at room temperature for ∼1 h) by Solid Phase Microextraction (SPME) inserting fibres consisting of three phases (divinyl benzene (DVB)/carboxen (CAR)/polydimethyl siloxane (PDMS), p/n 57328-U, Supelco, Sigma-Aldrich) into the headspace vials. Short-term to one-year storage in freezing conditions has been shown to have no significant impact on volatile components, and has been used to store biological samples, such as flowers and fruits, prior to subsequent volatile analysis (De Ancos *et al*., 2000; Qian *et al*., 2019). Fibres were kept in vials with thawed flowers for 1 hour of sampling, and were afterwards analysed using gas chromatography (HP6890, Hewlett-Packard, USA) coupled with a mass spectrometer (HP5973, Hewlett-Packard, USA) fitted either with a DB-5ms column (50 m × 0.2 mm ×0.33 µm, JandW Scientific, CA, USA) or a DB-wax column (polyethylene glycol, 30 m × 0.25 mm × 0.25 μm, JandW Scientific, CA, USA) using helium as carrier gas with a flow rate of 0.7 mL/min or 1 mL/min, respectively. Volatiles were desorbed from the fibres at 260°C in splitless mode (1 min). The oven was heated at a rate of 5°C/min from 40°C to 200°C and then at 10°C/min up to 260°C/min and held for 5 min. A scan range of m/z 45–400 and the ion source was set to 200°C, and the transfer line temperature to 260°C. Retention indices (RI) were calculated on both columns using the same linear gradient method with comparison to an n-hydrocarbon mixture (Sigma-Aldrich, St Louis, MO, USA, p/n 46827-U) using liquid injection in splitless mode (1 min sampling time). The main peaks in the total ion chromatogram of each sample were integrated using the HP Chemstation software, and the relative percentage abundance of peaks was determined. To characterise unknown compounds detected in two orchid samples, we used SPME combined with gas chromatography-high resolution mass spectrometry (GC-HRMS) recorded on a Waters GCT Premier TOF-MS using a DB-5ms column (30 m × 0.25 mm × 0.25 μm, JandW Scientific, USA). Method conditions were identical to above.

Scent data from the two orhids and the co-flowering plants were analysed by Principal Component Analysis (PCA), using R package *ade4*, version 1.7.18 and visualised using package *PCAviz*, version 0.3.37 (Dray et *al.*, 2007; Novembre and Dupanloup, 2019). Initial analyses revealed that the scent profile of *L. laevigatum* had major impact on the resolution of the PCA (Fig. S2), and we present the results of a PCA excluding this species. To evaluate the effect of species on the scent composition, we conducted a non-parametric multivariate analysis of variance (perMANOVA) using the function *adonis* with 999 permutations in the vegan R-package (Oksanen *et al*., 2013).

### Pollinator observations

In 2019, we conducted preliminary pollinator surveys across study sites, to select focal populations for later pollinator observations. We selected populations with as high fruit set as possible, where we expected high pollinator activity. We also identified the focal rewarding species for pollinator observations, by selecting the most common rewarding species across the orchid study populations.

#### Pollinator observations on Thelymitra

To identify pollinators of the two orchids and assess their behaviour, we conducted field observations during three flowering seasons (2020-2022; 11 - 26 August 2020; 13 - 20 August 2021; and 10 - 28 August 2022) in Tozer’s Bush Camp for *T. speciosa*, and in Waychinicup and Gull Rock National Park for *T. porphyrosticta*. Observations were made when temperatures were above 18°C (as measured with a Smartsensor AR827, set 20 cm above the ground) and flowers started to open. To identify pollinators of orchids and co-flowering plants, we used four GoPro Hero 5 cameras, placed 30 - 50 cm from the focal plants. Cameras were mounted on tripods at a height of ∼40 cm and set to record in medium-FOV (field of view) mode. Prior and post recording, we visually inspected flowers and noted the number of pollinia removed and deposited. A total of 195 15-min observation trials were conducted between 0930 and 1630 h, for a total of 121 hours and 87 min of observations. We recorded the number of visitors and their identity in each observation period. To increase the chance of pollinator visitation to orchids, we placed floral displays of common co-flowering rewarding plants (*Calectasia*, *A. caeruleum*, *D. hakeoides, M. suberosa*) nearby orchids (Scaccabarozzi *et al*., 2020). For each observation period, an array of co-flowering plants consisted of three glass vials separated by 10-20 cm and each containing 2-3 cut inflorescences of one of the rewarding species (with total 10–20 flowers). Vials were placed 0.5-1 m from flowering orchid individuals. Visiting insects were caught by net and the specimens were identified and vouchered at the Museum of Western Australia. Further, we identified all pollinators observed to remove and deposit orchid pollinia via visual examination of photos and video footage (Fig. 2). Because visitation rates were extremely low, we simply summed the number of visits for each species across all observation periods [**Supplementary Information, Table S3**]. We also searched for pre-existing records of insects caught visiting the orchids and/or co-flowering species in the collection at the Western Australian Museum [**Supplementary Information, Table S4**].

**Fig. 2.**
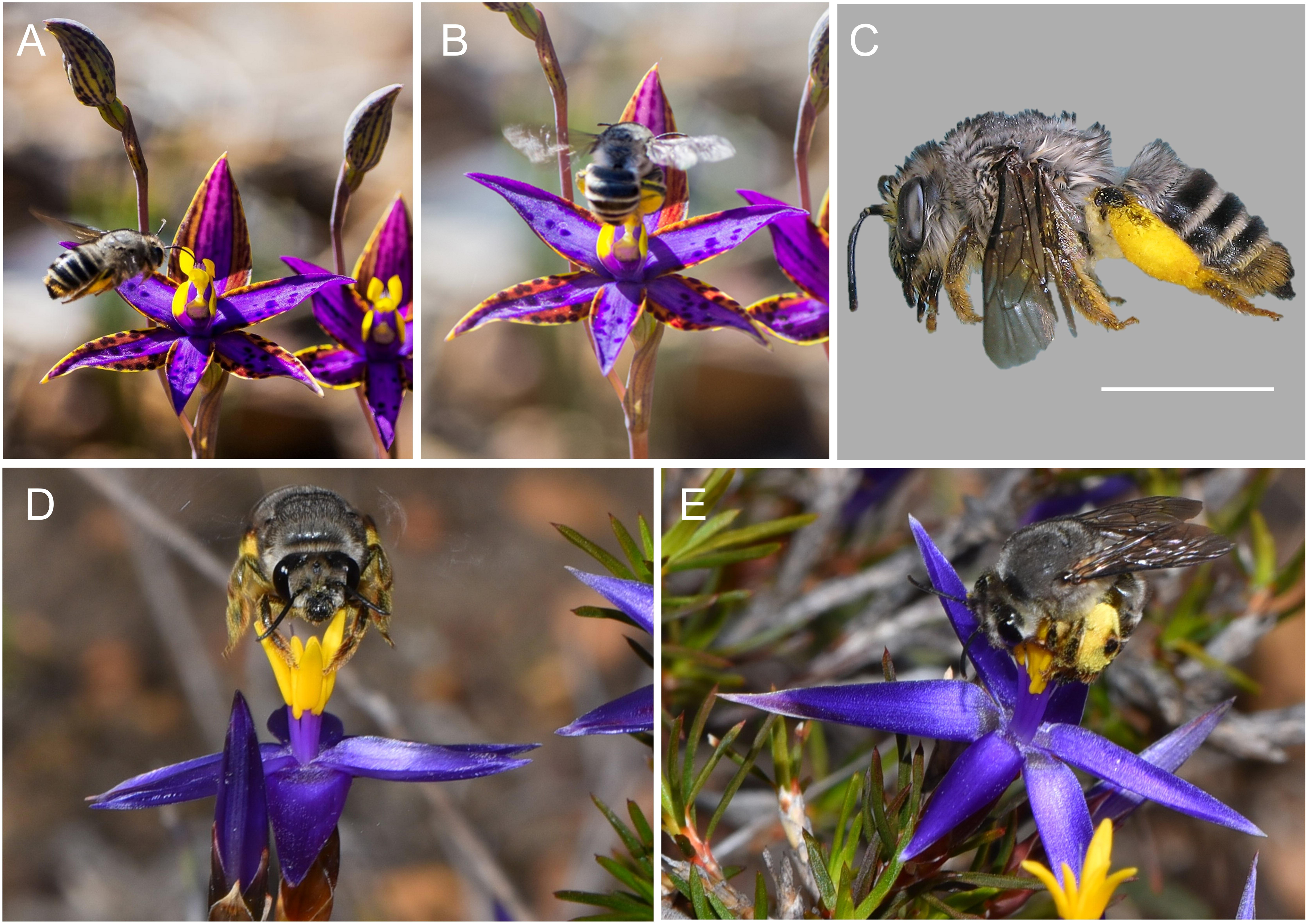
Bee pollinator **A:** approaching a *T. speciosa* flower, pointing antenna towards pseudo-anther; **B:** landing by curving its body around the pseudo-anther of a *T. speciosa* flower; and **C:** caught while buzzing on a tinsel lily, *Calectasia gracilis*. The behaviour in A and B is consistent with pollen foraging and buzzing. The individual bee has been identified as *Anthoglossa plumata* and carries tinsel lily pollen on scopa. **D-E:** *A. plumata* on tinsel lily flower, showing phases of buzz pollination. Scale bar: 0.5 cm. Photo credits: A-B: Christine Chester; C: Terry Houston; D-E: Kate Brown.

**Fig. 3.**
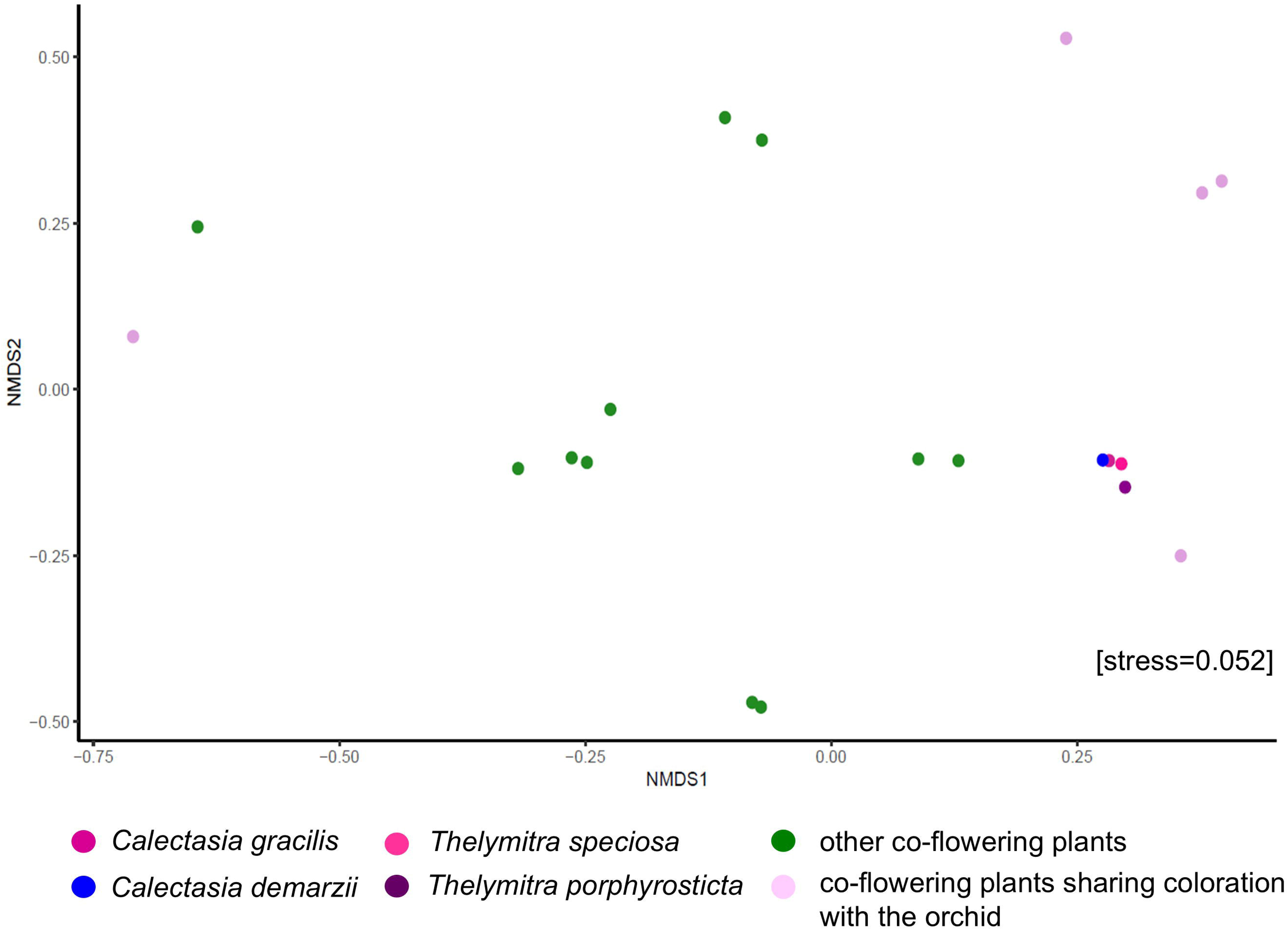
Non-metric multi-dimensional scaling plot based on plant and flower morphology (plant height, corolla width, corolla height, corolla symmetry, presence of exposed pollen, stamen/pseudoanthery height) of *Thelymitra porphyrosticta* and *T. speciosa*, *Calectasia demarzii* and *C. gracilis*, and 16 co-flowering species (see Table S2 for details).

To identify pollinator behaviour on orchids and on co-flowering plants, video material and stills were examined. Behaviour during visitation to a flower was divided into nectar versus pollen foraging. The following behavioural sequence was considered consistent with nectar foraging: approach to the flower by direct or zig-zag flight pointing head towards stigma, and landing on the tepal, attempting to forage nectar by pointing mouthparts towards the stigmatic area (Scaccabarozzi et al., 2020). Conversely, the following behavioural sequence was considered consistent with pollen foraging: approach to the flower by direct or zig-zag flight pointing antenna towards false anthers, and landing on the false anthers, and bending the body around the anthers (De Luca and Vallejo-Marín, 2013). Particularly, a behaviour where bees held anthers with their forelegs and display a recurved posture while attempting to manipulate them with their mandibles was considered suggestive of buzz pollination, where pollen is released from the anthers by vibrations of the thorax of the bee (Buchman, 1983; De Luca and Vallejo-Marín, 2013).

#### Pollinator observations on co-flowering plants

To determine if *Thelymitra* shares pollinators with the tinsel lilies and other common rewarding species, we conducted in-person pollinator observations at the same three study sites as above in the periods 14-26 August 2020, 15-20 August 2021, and 13 August-18 September 2022. Observations were made daily, between 1100 and 1500 h. On both *Calectasia* species, we performed a total of ten 5-min observation periods, yielding a total of 50 min observation. Again, we summed the number of visits for each species across all observation periods.

We used the same observation design on three other common rewarding species, that are potential pools of orchid pollinators, i.e., *A. caeruleum, D. hakeoides,* and *M. suberosa*. To also include some of the less common co-flowering species, we performed two 5-min observation periods for *Conospermum coerulescens*, *D. linearis*, *Hakea cucculata, H. lasiantha, H. psilocarpa, Isopogon formosus* and *P. squamata*.

### Orchid reproductive success in relation to the abundance of putative model species and rewarding community

To test if reproductive success of the orchid species was related to the abundance of the putative model species, the tinsel lilies (*Calectasia*), or to the total co-flowering rewarding community, we recorded pollinia removal and fruit production for all orchid individuals in 10×10 m plots in all study populations and years. For *T. speciosa*, we always included one plot per population, and for *T. porphyrosticta,* number of plots per population ranged from one to four, depending on orchid population size. Number of flowering orchids per plot ranged from three to 31 [**Supplementary Information, Table S5**]. For each orchid individual, we recorded total number of flowers, as well as the number of pollinia removed and the number of fruits produced. For each plot, we calculated pollinia removal and fruit production as the mean across individuals. In addition, we recorded number of flowers of *Calectasia* and other rewarding co-flowering species. Total number of flowers for each rewarding species and plot was estimated by counting number of inflorescences and multiplying by mean number of flowers per inflorescence, counted on five individuals. Total number of flowers was recorded as categories of ten (10=1-10 flowers, 20=11-20 flowers, 30=21-30 flowers, etc.). Across all populations and years, only one *T. speciosa* individual produced more than one fruit, and we treated fruit production as a binomial variable for this species. Neither mean pollen removal nor mean fruit production was related to the number of orchid individuals in the plot for any of the two species (all p>0.30, all r^2^<0.14).

To examine whether orchid reproductive success (number of pollinia removed and number of fruits produced per plant) varied among populations and years, we used a GLM with binomial distribution and logit link for *T. speciosa*, and GLM with Poisson distribution and log link with overdispersion correction for *T. porphyrosticta.* Both models included number of flowers as a covariate. To test whether orchid reproductive success was related to abundance of tinsel lilies and other rewarding species at the plot level, we used linear regressions and related mean pollen removal and mean fruit production per plot to number of i) *Calectasia* flowers (only *T. speciosa,* due to limited variation in abundance of tinsel lilies among *T. porphyrosticta* populations), ii) flowers with similar colour (*A. caerulea*, *A. simplex*, *C. caeruleum, D. linearis, M. suberosa*), and iii) total rewarding flowers, analysing each orchid species separately. In cases with a significant relationship between orchid reproductive success and abundance of flowers of similar colour, we also examined relationships for single rewarding species. In 2022, the tiny TZ3 population (n=3) of *T. speciosa* had unusually high fruit production, and we conducted regressions both including and excluding this outlier. It should be noted that in these analyses, data points (plot means) cannot be considered fully independent, as some populations of *T. porphyrosticta* contained more than one plot, and some populations and plots were included multiple years. Analyses were performed using JMP 17.0.0 (JMP®, Statistical Discovery, 2022 - SAS Institute Inc., Cary, NC, 1989–2023).

## RESULTS

### Floral trait similarity between orchids, tinsel lilies, and other rewarding species

#### Morphology

The NMDS plot indicated that orchids and tinsel lilies were more similar to each other than to the rest of the co-flowering plant community [**Supplementary Information, Fig. 3**]. The flowers of *Thelymitra* were considerably fewer and larger than those of *Calectasia* [**Supplementary Information, Table S6**]. The perMANOVA analysis revealed a significant effect of plant species, indicating that scent profiles varied significantly among species (F = 12.11, df = 10, r² = 0.7, p = 0.001).

#### Colour

Colour analysis using the hexagon bee vision model showed that the average flower colour loci of the orchids, tinsel lilies, and other rewarding plants with similar flower colour, all corresponded to the bee-subjective UV-blue region of the colour hexagon (Fig. 4A). The two species with deviating flower colour fell in the blue-green and ultraviolet regions of the hexagon (Fig. 4A). *Thelymitra speciosa* was most similar to its co-flowering tinsel lily (mean colour distance to *C. gracilis:* = 0.03 hexagon units), while *T. porphyrosticta* was more similar to *A. caerulea* (0.03) and *C. caeruleum* (0.04) than to *C. demarzii* (0.09) (Fig. 4A). Colour loci coordinates for individuals of the two orchids overlapped, and loci for *T. speciosa* also overlapped with individuals of *C. gracilis* and *A. caerulea*, while loci for *T. porphyrostricta* only overlapped with *A. caerulea* (Fig. 4A). False colour photos in bee view showed that flowers of *T. speciosa* closely match the colour of the tinsel lily *C. gracilis*, regarding both the star-shaped petals and the false anthers (Fig. 4B).

**Fig. 4.**
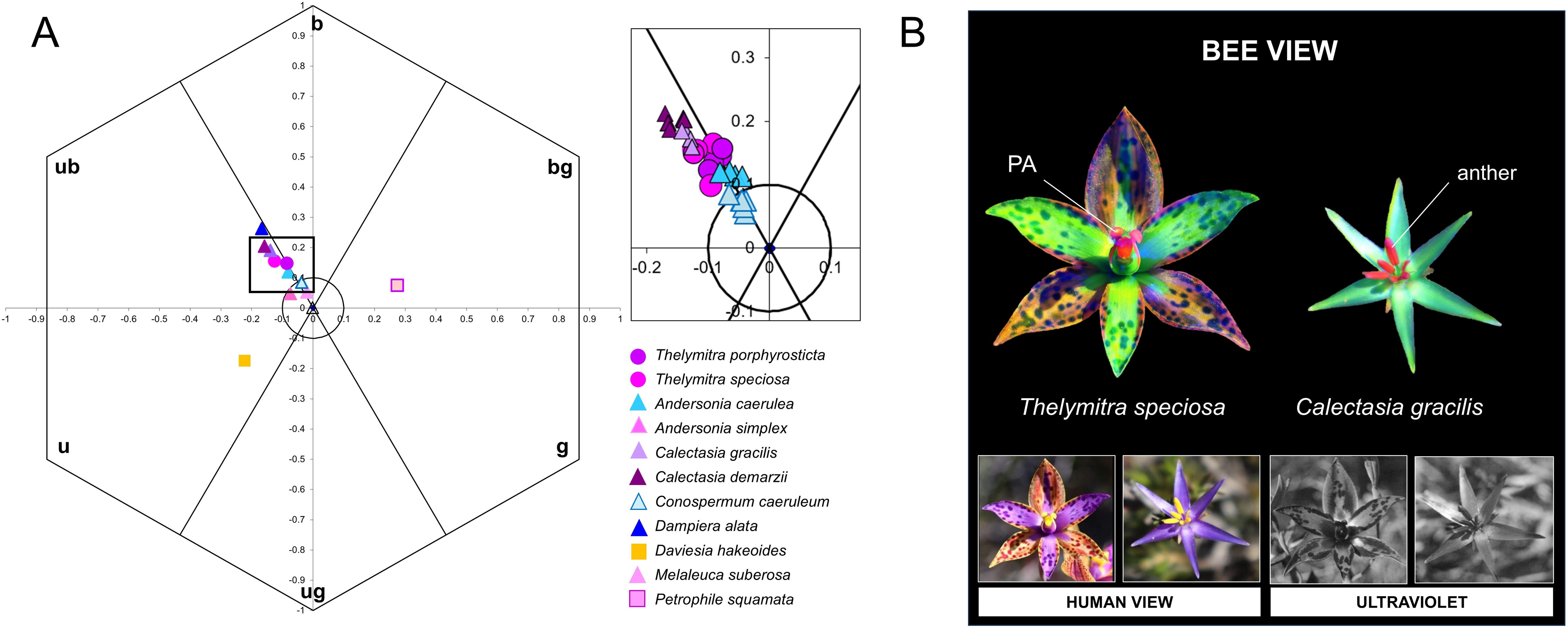
**A**: Mean flower colour loci on tepals of the deceptive orchids *Thelymitra porphyrosticta* and *T. speciosa*, the co-flowering rewarding tinsel lilies *Calectasia demarzii* and *C. gracilis*, and five other co-flowering rewarding species with similar flower colour; *Andersonia caerulea, A. simplex*, *Conospermum caeruleum, Dampiera linearis,* and *Melaleuca suberosa.* Two co-flowering species with deviating flower colour, *Petrophile squamata* and *Daviesia hakeoides,* are included for comparison. Inset shows individual colour loci of the orchids, tinsel lilies, and *A. caerulea* and *C. caeruleum* (n=6 per species). Calculations were based on the hexagon colour model of bee vision (u=ultraviolet, ub=ultraviolet-blue, b=blue, bg=blue-green; g=green; ug=ultraviolet-green; Chittka, 1992). **B**: False colour photography of *T. speciosa* and *C. gracilis* in bee view. Flowers are photographed in visible light (human view), UV, and transformed in false colour photography (bee view); Verhoeven *et al*., 2018.

#### Scent

In *T. speciosa,* the GC-MS analysis identified 10 major compounds (above 1% of relative abundance) that were all shared with its co-flowering tinsel lily *C. gracilis* [Fig. 5; **Supplementary Information, Table S6**]. Scent was dominated by limonene, that ranged in concentration from 21.1% to 25.9%, followed by C_11_H_18_O_2_ (7.9%-33.70%), CH_15_H_30_ (0%-16.8%), *p*-Cymene (4.3%-5.8%), γ-Terpinene (3.5%1-3.9%), and other compounds at lower concentrations. In *T. porphyrosticta*, we identified four compounds, that all were shared with *T. speciosa* (Fig. 5, Table 6). Samples of *C. demarzii* contained only C_11_H_18_O_2_ (2.2%-17.9%), that was also present in both orchid species and in *C. gracilis* (Fig. 5). Some orchid compounds were also present in samples of other species in the rewarding community (*A. caerulea:* limonene, *β*-Pinene; *M. suberosa:* limonene, eucalyptol; *L. laevigatum*: *p*-Cymene), but most orchid compounds were absent from species other than tinsel lilies (Fig. 5). The PCA showed that scent profiles of *T. speciosa* samples overlapped with those of *C. gracilis* (Fig. 6A), with similarity driven by multiple compounds (Fig. 6B). Scent profiles of *T. porphyrosticta* samples were close to both *Calectasia* species (Fig. 6).

**Fig. 5.**
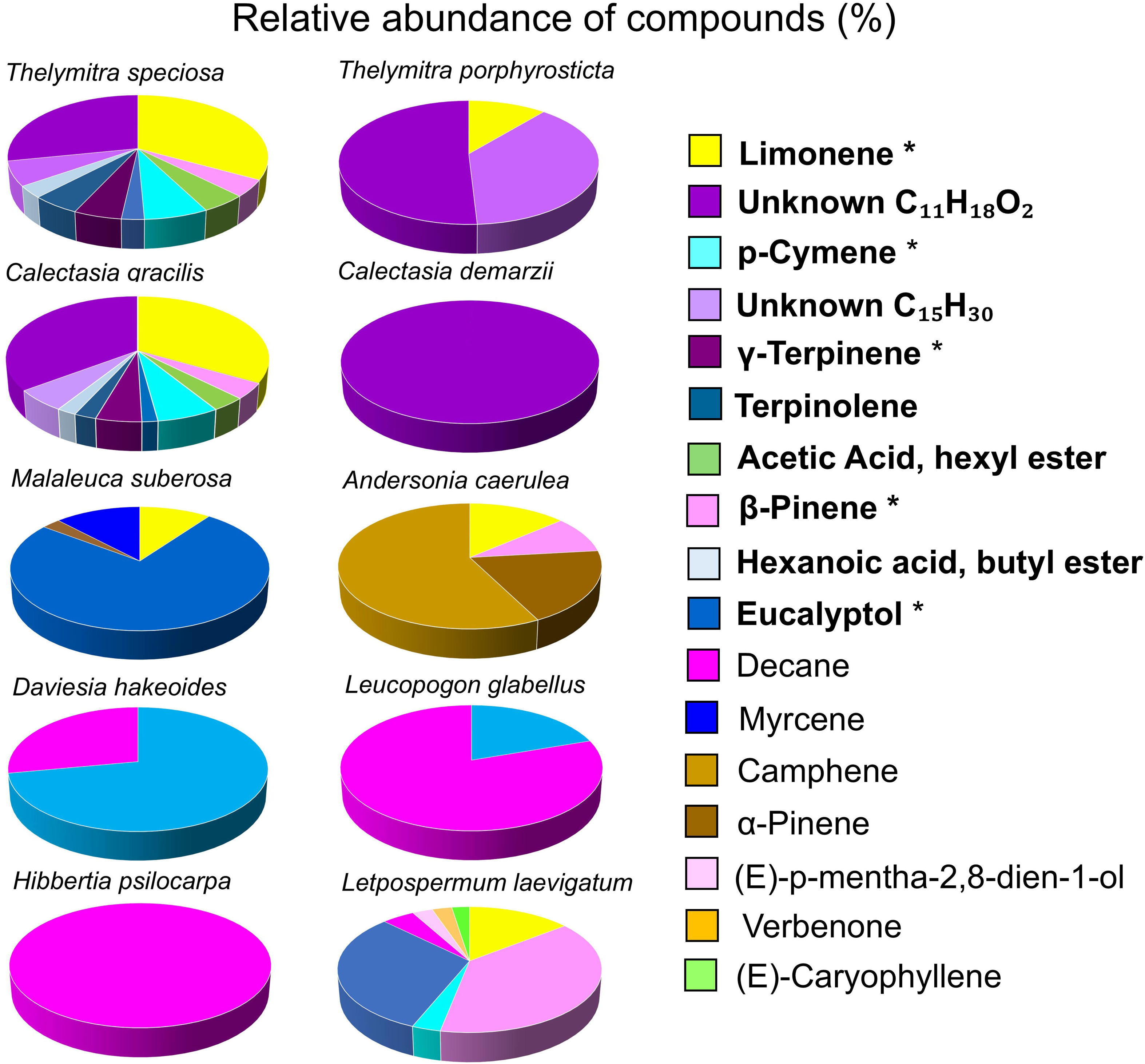
Relative abundance of flower scent compounds (%) identified by GC-MS analysis of samples from the two orchids *Thelymitra porphyrosticta* and *T. speciosa*, the two tinsel lilies *Calectasia demarzii* and *C. gracilis*, and six other co-flowering rewarding species. Compounds above 1% relative abundance in samples are included. Compounds detected in the orchids are indicated in bold. *Confirmed with commercial standard.

**Fig. 6.**
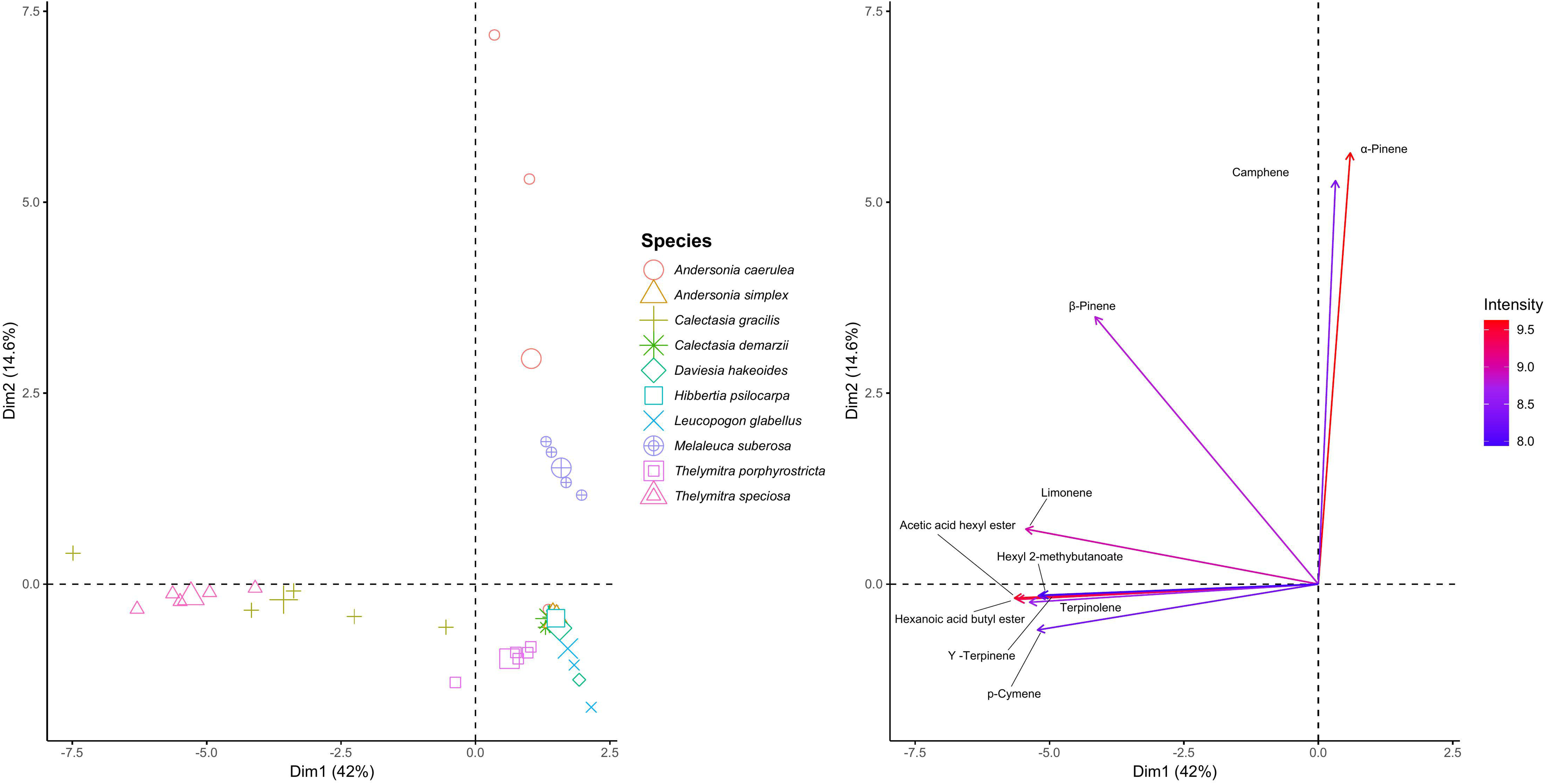
Principal Component Analysis (PCA) of floral scent of orchids and co-flowering plants. **A**) Biplot representing scent samples from the orchids (*Thelymitra porphyrosticta* and *T. speciosa*), the tinsel lilies (*Calectasia demarzii* and *C. gracilis*), and six other co-flowering rewarding species. **B**) Biplot showing the compounds associated with the PCA axes 1 and 2. The colour variation of the arrows represents the intensity of each compound’s contribution to the distribution of scent samples in the biplot shown in Fig. 6A.

### Pollinator observations on *Thelymitra* and co-flowering plants

Both orchid species shared bee pollinators with the tinsel lilies and other rewarding plants, and the pollinators showed a behaviour associated with buzz pollination on both orchids and tinsel lilies. A total of 19 visits were observed on the two orchids, of which 15 were by bees, three by beetles, and one by an ant [**Supplementary Information,** Table S3]. Of the 15 bee visits, five were by *Anthoglossa plumate* females, five by *Apis mellifera,* three by *Leioproctus* sp., one by *Lasioglossum* (*Parasphecodes*) sp., and one by *Trichocolletes erythrurus.* The video footage captured an *A. plumata* bearing pollinia at the tip of the abdomen, depositing traces of pollinia on three flowers of *T. porphyrosticta* and extracting the pollinia of a flower on the same individual orchid [**Supplementary Information -** Video & Fig. S3]. Another *A. plumata* individual was observed to remove pollinia from *T. speciosa* and also from another individual of *T. porphyrosticta*. The bee afterwards moved to *A. caerulea* for nectar foraging. Further records showed two *Lasioglossum* sp. bees on the orchid flowers. In addition, two *A. plumata* bees were observed to carry orchid pollinia while foraging on *Hakea lasiantha* and *C. gracilis*. *A. plumata* bees visiting orchids displayed a behavioural sequence suggestive of buzz pollination. Bee visits lasted less than one second. One *Lasioglossum* and one *Leioproctus* individual attempted to forage nectar on the orchids. Similarly, *A. mellifera* demonstrated typical nectar foraging behaviour.

Based on observations of contact with the reproductive structures, the tinsel lilies *C. gracilis* and *C. demarzii* were pollinated by *Leioproctus* sp. and *A. plumata* [**Supplementary Information,** Table S3]. Bees of both species sonicated while contacting anthers of the tinsel lilies, releasing pollen. These visits lasted more than two seconds.

Observations on other co-flowering plants showed that the pollinators of orchids and tinsel lilies also visited other species in the community (see Table S3 for details), and all visiting bees exhibited nectar foraging behaviour. A total of 24 floral visitors were caught during observations on orchids and co-flowering plants. Nine *Anthoglossa* bees caught on target plants in the study sites were found in pre-existing records at the Western Australian Museum [**Supplementary Information,** Table S4].

### Orchid reproductive success in relation to abundance of tinsel lilies and other rewarding species

In *T. speciosa*, pollen removal varied significantly among populations (χ^2^ _3,117_=21.4, p<0.001, range=0-0.91, range=0.11-0.18, n=124) and years (χ^2^_2,117_=12.9, p=0.016, range=0.15-0.33), but fruit production did not (all p>0.05, population range=0.05-0.10, year range=0.03-0.19). In plots, mean number of fruits increased with the abundance of both *C. gracilis* (β=0.0040, r^2^=0.57, F_1,9_=10.6, p=0.012, n=10, Fig. 7A), and flowers of similar colour (β=0.0017, r^2^=0.64, F_1,9_=14.5, p=0.0052, Fig. 7B) when the outlier population was included, but none of the relationships were statistically significant when this population was excluded (β=0.0080, r^2^=0.03, F_1,8_=0.21, p=0.66 and β=0.0059, r^2^=0.09, F_1,8_=0.75, p=0.42, respectively) (Fig. S4).

**Fig. 7.**
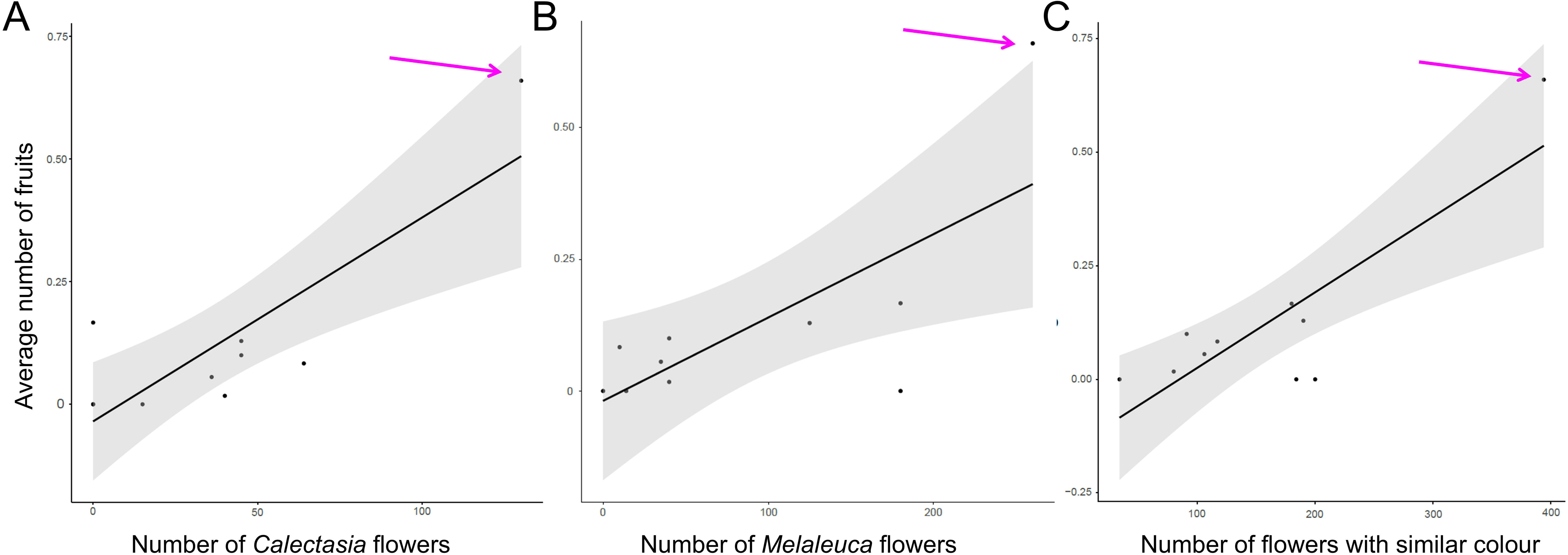
Mean number of fruits in 10 plots of *Thelymitra speciosa* in relation to total number of flowers of **A**) *Calectasia gracilis,* **B**) *Melaleuca suberosa,* and **C**) rewarding species sharing flower colour with the orchids. The arrow in each panel indicates the outlier driving the relationship.

The relationship with flowers of similar colour when including the outlier was mainly driven by the abundance of *M. suberosa* (β= 0.0017, r^2^=0.56, F_1,9_=9.99 p=0.013, Fig. 7C). Mean pollen removal in plots was not significantly related to the abundance of tinsel lilies or other co-flowering plants (all p>0.28).

In *T. porphyrosticta*, pollen removal varied significantly among populations (χ^2^_4,149_=11.5, p=0.021, range=0.09-0.47, n=156) and years (χ^2^_1,149_= 12.4, p=0.004, range=0.16-0.45), but fruit production did not (all p>0.05, population range=0-0.41, year range=0.22-0.37). In plots, neither pollen removal nor fruit production was significantly related to any component of the rewarding community (all p>0.05, n=16).

## DISCUSSION

In this study, we investigated whether pollination of the deceptive Queen of Sheba orchids, *T. speciosa* and *T. porphyrosticta*, occurs via finely tuned mimicry of tinsel lilies (*Calectasia* spp.), or alternatively through a form of generalized food deception. We found that *T. speciosa* is nearly indistinguishable from its co-flowering tinsel lily *C. gracilis* regarding both flower colour and scent, and that *T. porphyrosticta* also resembles its co-flowering species *C. demarzii*, although with less precision. The closest colour match to *T. porphyrosticta* was found in *A. caerulea* and though its major scent compound was shared with *C. demarzii*, it also contained additional compounds. Orchids and tinsel lilies also shared pollinators, and the main visitor on orchids displayed a behaviour associated with buzz-pollination of tinsel lilies. Similarity in flower traits and pollination indicate specific mimicry towards tinsel lilies in both species, but the lack of any relationship between model species abundance and orchid fitness raises the question of how efficient this strategy is.

### Similarity of floral traits in orchids, tinsel lilies and other rewarding species

The two orchids had overlapping flower colour, but only *T. speciosa* closely matched the colour of its co-flowering tinsel lily, *C. gracilis* (Fig. 4A). Euclidean distances between colour loci of single individuals of the two species were always less than 0.06 hexagon units, which represents the minimum distance where honeybees can distinguish colours (Dyer and Chittka, 2004 a,b; Giurfa, 2004). The false colour photography in bee view also documented striking similarity of both anther contrast and petal colour between *T. speciosa* and *C. gracilis*, suggesting that bees may perceive these flowers as identical (Fig. 4B). The distances between colour loci of individuals of *T. porphyrosticta* and *C. demarzii* were larger (range 0.07-0.11), suggesting that bees are able to separate the colour of these two species. Still, the accuracy of colour vision models can depend on light conditions and background colour (Bukovac et al. 2017; Finnell and Koski 2021), and we do not know if the colour sensitivity used in the honeybee visual model precisely match those of the native bees visiting Queen of Sheba orchids. Moreover, the results indicated that orchids were morphologically more similar to tinsel lilies than to the remainder of the co-flowering community. The orchid flower resembles a larger version of the tinsel lily flower, displaying a supernormal stimulus, a more salient stimulus compared to model plants (Leonard *et al*., 2011; Scaccabarozzi *et al*., 2023). This stimulus does not violate the mimicry but might increase further floral attractiveness. It is possible that, together with shared flower morphology and conspicuous yellow anthers, colour similarity of *T. porphyrosticta* and *C. demarzii* is sufficient to confuse the pollinators. Interestingly, flower colour of *T. porphyrosticta* was most similar to co-flowering *A. caerulea* and *C. caeruleum*, with mean distances between colour loci of 0.03 and 0.04, respectively. These two species are not morphologically similar to the orchid, they are not buzz pollinated and have no contrasting yellow anthers. The existence of a specific model species does not exclude the possibility that other rewarding species may contribute to pollinator attraction, and taken together, the results suggests that both orchids should benefit from colour resemblance to co-flowering *Calectasia*, but also from similarity to more common species in the community. A parallel pattern emerged from the analysis of scent. The fragrances of *T. speciosa* and *C. gracilis* were remarkably similar, both in terms of compounds present and their relative proportions [Fig. 5; **Supplementary Information,** Table S6]. The two species shared a complex scent profile including ten compounds. In contrast, *T. porphyrosticta* had only one compound in common with *C. demarzii*. The shared, unknown compound was the only one present in the tinsel lily and was also the dominant one in the orchid. In addition, both orchid species shared a few compounds with other co-flowering plants, such as *M. suberosa* and *A. caerulea*. Overall, scent compounds produced by the orchids were present in their mimics but rare or absent in other co-flowering species, supporting the hypothesis of finely tuned mimicry. Although scent is known to play a fundamental role in sexually deceptive orchids, less is known about its role in food mimicry (Schlüter and Schiestl, 2008). Choice experiments using artificial flowers in a bee-pollinated food mimicry system found that discrimination of the mimic was poorest if it shared the same scent as the model (Gumber and Kunze, 2001), indicating that the optimal strategy for a mimic should be to replicate the scent of its specific model species. In a more generalised situation with multiple models, having weak or no scent may be better than mimicking only one model (Gumber and Kunze, 2001). As such, the precise scent match between *T. speciosa* and *C. gracilis* strongly indicates that the orchid mimics the tinsel lily, with scent reinforcing visual similarity. In sum, our findings indicate that scent can be a key trait involved in food mimicry, but trait manipulations will be necessary to clarify the role of visual and scent signals in this system.

The floral bouquet of *T. speciosa* contained several compounds that frequently occur in the scent of bee-pollinated orchids, such as limonene, *β*-pinene, and *p*-cymene (Cancino and Damon, 2007; Watteyn *et al*., 2021; Watteyn *et al*., 2023). Common constituents of scents of perfume flowers, such as eucalyptol and terpinolene (Zito *et al*., 2013; Milet-Pinheiro *et al*., 2022) were minor components of the orchid scent. While scent profiles can be very complex, specific key chemicals may be the ones that trigger bee responses (Dötterl *et al*., 2005), and a bioassay would be necessary to confirm the role of individual scent components in floral mimicry. Former studies have shown that bees can rely on pollen odours to detect host plants (Dobson and Bergström, 2000), and that buzz-pollinating bees can prefer feeding anthers over pollinating anthers, due to different volatiles in the two anther types (Solís-Montero *et al*. 2018). It would be interesting to test if false anthers in the orchids emit volatiles that attract visitors or elicit the buzzing behaviour of bees.

Tinsel lilies comprise a range of species with similar flower colour and shape (George, 1986; Barrett and Dixon, 2001), and although we only found a single species together with each orchid during this study, more than one species could co-occur with orchid populations over time. Considering this, an imperfect mimicry rather than an exact one may increase the chances of relying on multiple, more broadly similar existing models. Selection for convergence can be expected to be stronger on the most abundant model species, and it is possible that differences in the precision of phenotypic matching between the two orchids reflect differences in long-term stability of the model-mimic association. Detailed information on the historic distribution of orchids and tinsel lilies would be necessary to evaluate this.

### Pollinator observations on *Thelymitra* and co-flowering plants

Pollinator sharing between models and mimics is a crucial criterion for evaluating the occurrence and effectiveness of floral mimicry (Roy and Widmer, 1999; Johnson and Schiestl, 2016). Due to the very low visitation rates experienced by the orchids, we recorded only a few pollinator visits during two years of observations. However, on both orchid species, *A. plumata* bees removed and deposited orchid pollinia traces [**Supplementary Information-Video**], and this bee species also visited the tinsel lilies. *A. plumata* bees also displayed typical buzz-pollination behaviour on *Calectasia*, and behaved in a similar way on the orchids, attempting to seek pollen rewards. Tinsel lilies were the only buzz-pollinated plants present among the co-flowering species, suggesting that the observed behaviour of *A. plumata* on orchids could represent a conditioned response due to pollinator learning on *Calectasia*. However, the behaviour could also be due to an unconditioned response to anther-like structures resembling those of buzz-pollinated plants (Russel *et al*. 2017). We also identified other bees acting as potential orchid pollinators, such as *Lasioglossum* (Parasphecodes) sp. on *T. porphyrosticta,* and *Leioproctus* sp. on *T. speciosa*. These species attempted to forage for nectar, directing the head towards the stigmatic surface of the orchid flowers. This behaviour could lead to pollen transfer by placing pollinia on the bee’s head, which it then deposits during a subsequent visit to a flower. Both taxa also foraged for nectar on *Andersonia* and *Melaleuca*. Due to the scarcity of tinsel lilies in the habitat, the involvement of a range of polylectic bees displaying both pollen and nectar-seeking behaviour may benefit the orchids. Quantifying pollen transfer efficiency by bees displaying pollen-versus nectar-feeding behaviour could clarify their role in the pollination ecology of the orchids.

### Orchid reproductive success relative to abundance of tinsel lilies and other rewarding plants

If Queen of Sheba orchids specifically resemble tinsel lily flowers, we would expect orchid reproductive success to be greater when the model is more abundant (Anderson and Johnson, 2006). However, results did not convincingly support that male or female reproductive success increased with abundance of tinsel lilies. Although mean fruit production of *T. speciosa* increased with number of *C. gracilis* flowers, a similar relationship was found with the total number of rewarding flowers of similar colour, and both relationships were driven by a single orchid population, consisting of three flowering individuals with high fruit set. In *T. porphyrosticta*, reproductive success was not related to any component of the rewarding community. These results contradict expectations regarding both specific mimicry and generalised food deception. Both low orchid pollination success and limited variation in abundance of tinsel lilies make it difficult to test relationships, and in future studies, manipulations of model and magnet abundance could provide a clearer answer.

The group of Queen of Sheba orchids is of increasing conservation concern (Florabase, 1998). Habitat preservation and pollinator requirements (food plants) are key aspects in preserving orchids, whether they are existing or reintroduced. Knowledge of the pollination strategy can guide conservation efforts, and although we found weak relationships between orchid reproductive success and the rewarding community, the current results suggest that conservation practices focused on *T. speciosa* and *T. porphyrosticta* should promote co-flowering tinsel lilies, but also a habitat with mixed floral resources, especially of species sharing flower colour with the orchids.

### Conclusions

In summary, the results suggest that the two Queen of Sheba orchids specifically mimic co-flowering tinsel lilies. The striking phenotypic similarity of *T. speciosa* and *C. gracilis* involves multiple floral traits and is consistent with a situation where the orchid has experienced stable selection to mimic the model. Although less striking, *T. porphyrosticta* also resembles its co-flowering tinsel lily, *C. demarzii,* more than any other co-flowering species. However, these findings contrast with the lack of evidence that resemblance is adaptive, and it remains to be shown to what extent orchid reproductive success is influenced by the abundance of tinsel lilies and other rewarding species. Overall, the *T. variegata* complex offers an interesting system for studying the evolution of specific mimicry versus generalised food deception.

## SUPPLEMENTARY INFORMATION

Supplementary information is available online at www.aob.oxfordjournals.org and consist of the following. Fig. S1: Most common co-flowering species that to the human eye were similar in colour to *Thelymitra* species; Fig. S2: Principal Component Analysis (PCA) of floral scent of orchids and co-flowering plants; Fig. S3: Stills from the video showing bee pollinating a flower of *Thelymitra*; Fig. S4: Mean number of fruits in 10 plots of *Thelymitra speciosa* in relation to total number of flowers. Table S1: Detailed habitat description of all sites with study populations; Table S2: Floral traits of *Thelymitra* spp., *Calectasia* spp. and the most abundant co-flowering species; Table S3: Observations of floral visitors on the two orchid species and tinsel lilies; Table S4: List of insects caught on *Thelymitra* spp. and on co-occurring plants; Table S5: List of the plots of *Thelimitra speciosa* and *T. porphyrosticta* populations; Table S6: Compounds with compound class present in the orchids *Thelymitra speciosa* and *T. porphyrosticta,* and in co-flowering plant samples; Video: showing *Anthoglossa* behaviour associated with buzz pollination on flowers of *Thelymitra*. All the video imagery presented is in slow motion.

## ACKNOWLEDGEMENTS

We thank Frank Busetti, Stefan Dötterl, and Zong-Xin Ren for assistance and advice on chemical analysis; Max Massi, Liz Barbour, and Peter Chapman for equipment and facility support; Russell Barrett, Andrew Brown, Paul Armstrong, and Sarah Barrett for help with plant identification and locating orchid populations; Stig Dalström and Darryl Saffer for research support and advice; Katharina Nargar for contributing to manuscript discussions; Rob Tozer for logistical support; and Christine Chester, Mick Hurds, Lisa Wilson, and Kate Brown for photography and site location assistance. This work was supported by private donations received during the screenings of the documentary *Search for the Queen of Sheba* by Darryl Saffer and Stig Dalström, held in the City of Albany, Shire of Mundaring, Shire of Augusta-Margaret River, Shire of Collie, Bunbury Orchid Society, and Shire of Ravensthorpe. We extend our sincere thanks to all for supporting our research and outreach activities.

---

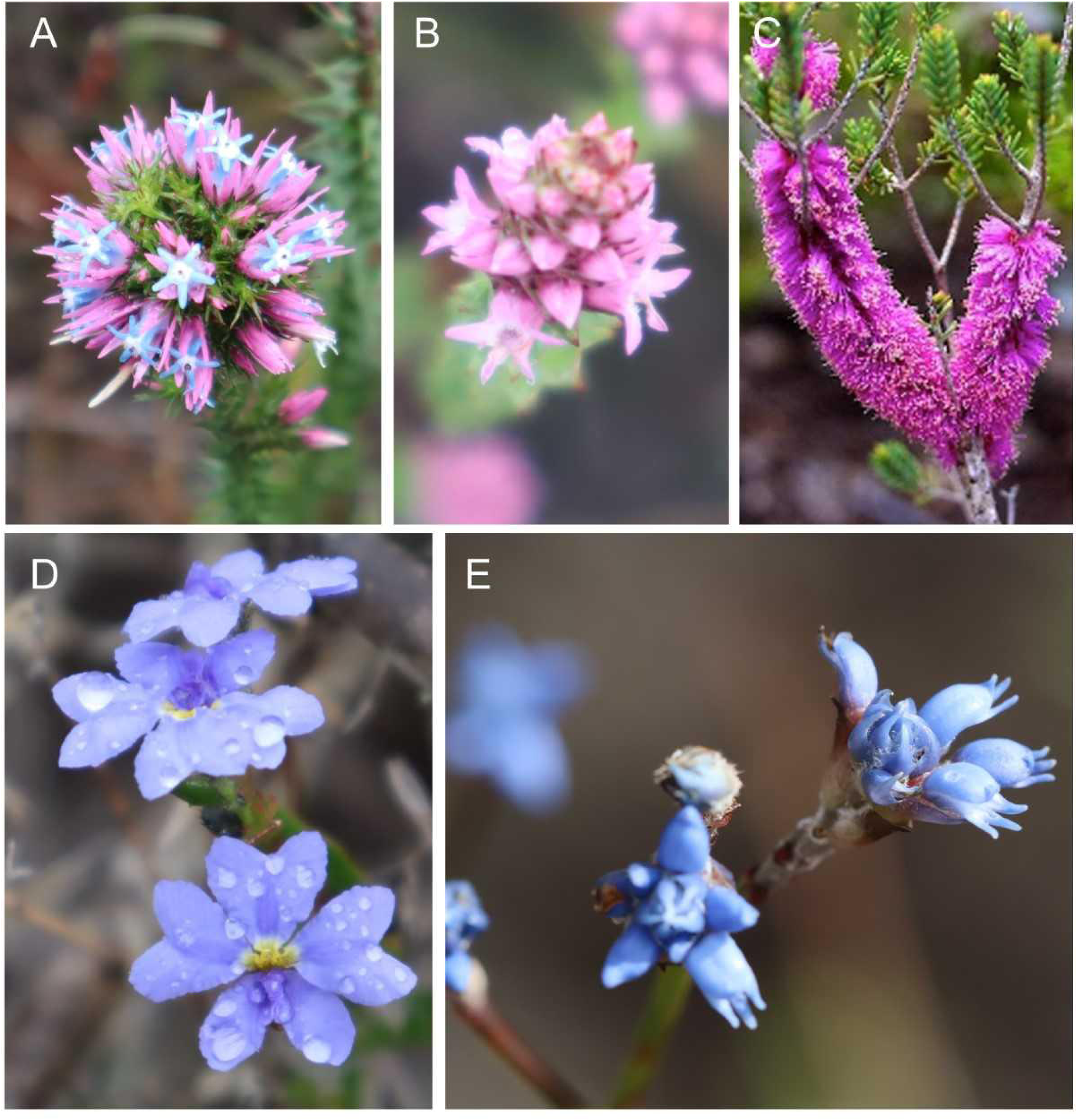

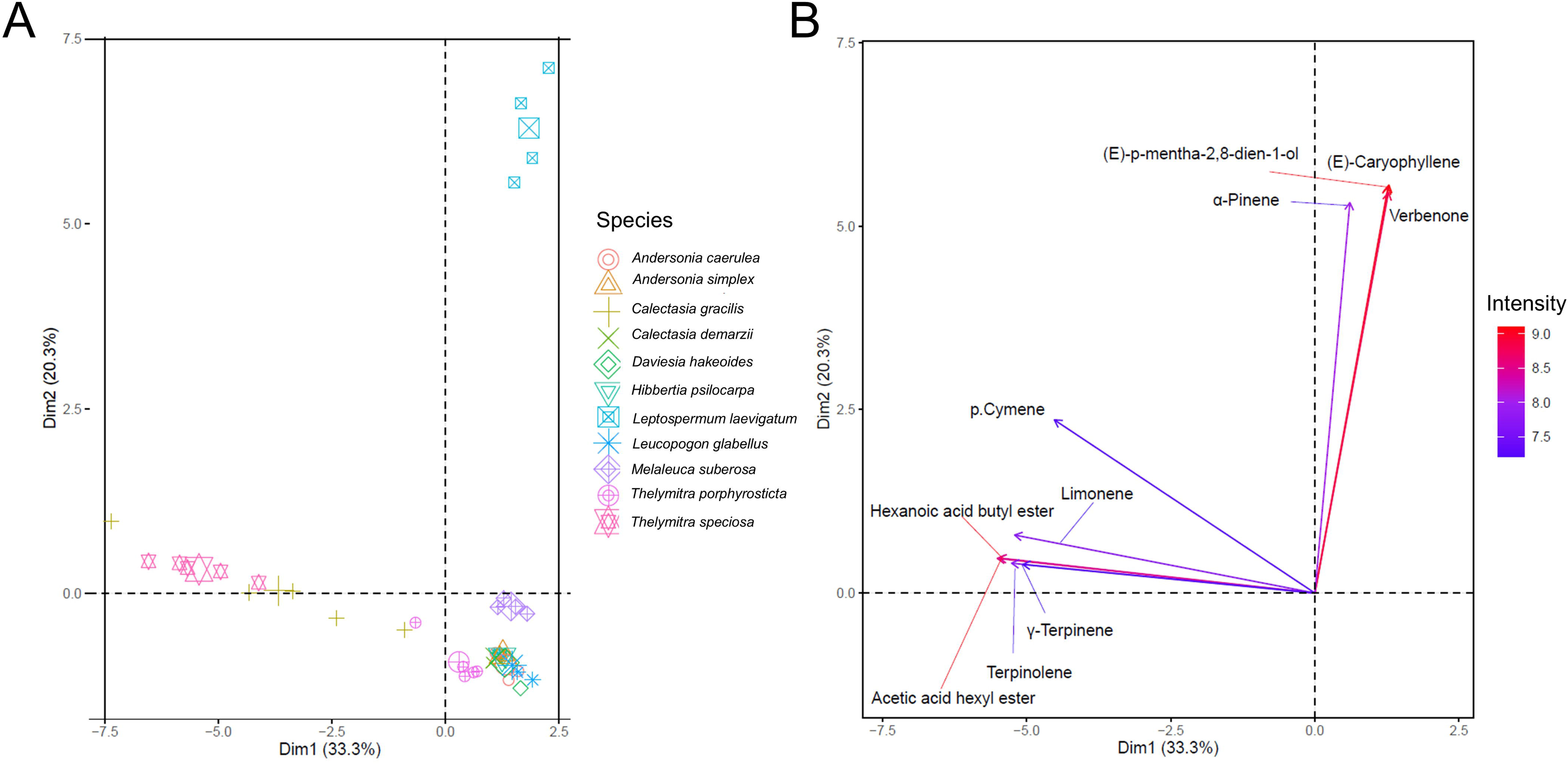

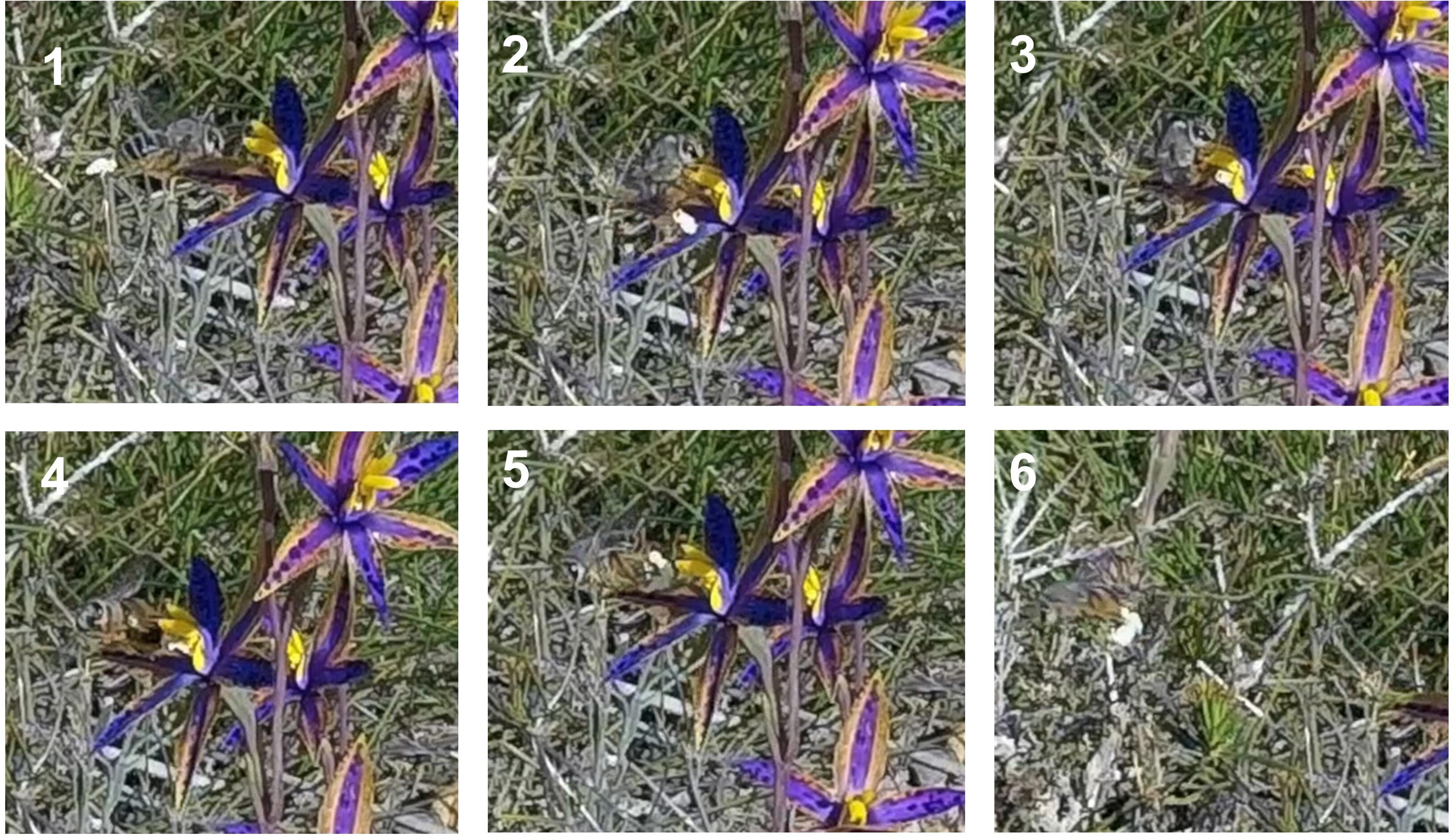

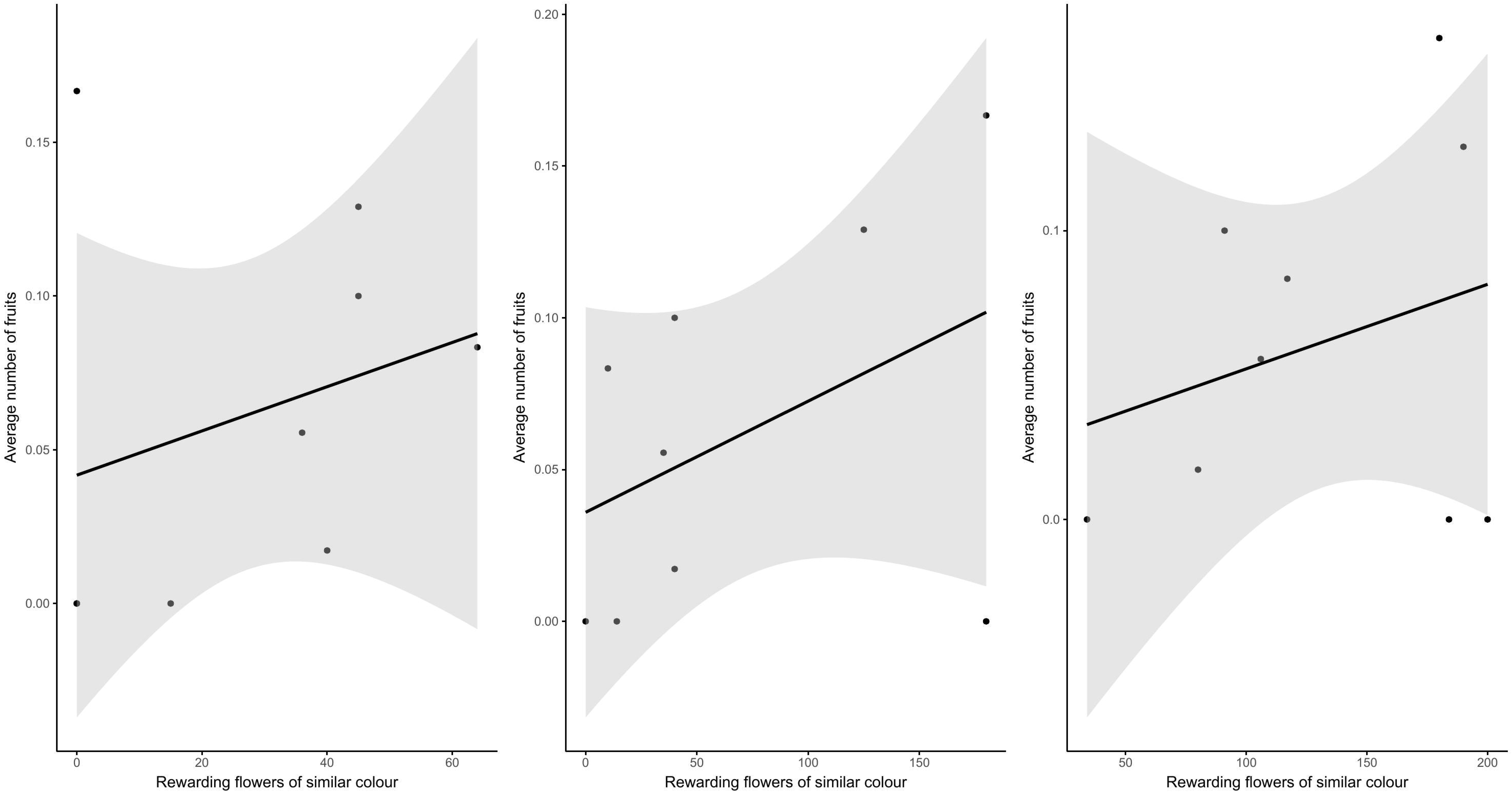

## Notes

### Competing Interest Statement

The authors have declared no competing interest.

