## Supplementary material for "Nuances of floral mimicry in Queen of Sheba orchids"

**SUPPLEMENTARY INFORMATION, Figure captions**

**Figure S1:** Common nectar-producing species that were co-flowering with the *Thelymitra* species, and that to the human eye were similar in flower colour.

**Figure S2:** Principal Component Analysis (PCA) of floral scent of orchids and co-flowering plants. **A**) Biplot representing scent samples from the orchids (*Thelymitra porphyrosticta* and *T. speciosa*), the tinsel lilies (*Calectasia demarzii* and *C. gracilis*), and seven other co-flowering rewarding species. **B**) Biplot showing the compounds associated with the PCA axes 1 and 2. The colour variation of the arrows represents the intensity of each compound's contribution to the distribution of scent samples in the biplot shown in Fig. 6A.

**Figure S3:** Stills from the video showing bee pollinating a flower of *Thelymitra*. 1: An *Anthoglossa* bee approaches a *Thelymitra* flower, pointing its antennae toward the pseudoanthers while carrying orchid pollinia. 2: The bee begins to turn its body, curving its abdomen over the pseudoanthers. 3: The bee pollinates the flower by depositing part of the pollinia onto the stigma. 4: The bee extracts a new pollinium from the flower (see the viscidium attaching to the bee abdomen). 5: The bee moves away from the flower, now carrying a visibly larger pollen mass (the previously held pollinia plus the newly collected one). 6: The bee flies away, carrying a large pollen mass.

**Figure S4:** Mean number of fruits in 9 plots (outlier removed) of *Thelymitra speciosa* in relation to total number of flowers.

**SUPPLEMENTARY INFORMATION, Table S1-S6**

**Table S1**. Detailed habitat description of all sites with study populations of *Thelymitra prophyrosticta* and *T. speciosa*, surveyed between 2020 and 2022.

| Site | Species | Vegetation description | Orchid populations | Co-flowering species |
|---|---|---|---|---|
| Tozer bushacamp site | *Thelymitra speciosa* | Dense *Malaleuca* dominant shrublands with open areas of low mixed vegetation communities. The upper canopy vegetation comprises Malle Eucalypt species and *Eucalyptus pleaurocarpa* being the dominant species. | TZ1, TZ2, TZ3, TZ4 | *Acacia* species, *Andersonia caerulea, Calectasia gracilis, Dampiera linearis, Daviesia hakeoides, Grevillea, Hakea obliqua, Hibbertia psilocarpa, Leucopogon glabellus, Malaleuca suberosa.* |
| Waychinicup National Park | *Thelymitra porphyrosticta* | Dense shrubland dominated by *Banksia, Hakea* and *Petrophile*. The upper canopy vegetation comprises scattered *Eucalyptus marginata* and *Nuytsia floribunda.* | WAY3 | *Acacia browniana, Andersonia caerulea, Banksia baxteri, Banksia coccinea, Calectasia demarzii, Conospermum caeruleum, Dampiera linearis, Hakea cucculata, Hakea lasiantha, Hakea trifurcata, Hibbertia psilocarpa, Isopogon formosus, Malaleuca striata, Petrophile squamata, Leucopogon glabellus.* |
| Gull rock National Park | *Thelymitra porphyrosticta* | *Banksia* woodland mixed with she-oak forest. | GR4, GR5, GR6, GR7 | *Acacia* species, *Agonis marginata, Allocasuarina fraseriana, Andersonia caerulea, Andersonia simplex, Banksia coccinea, Calectasia demarzii, Conospermum caeruleum, Dampiera linaris, Daviesia hakeoides, Dryandra formosa, Hakea elliptica, Leucopogon glabellus, Petrophile squamata.* |

**Table S2.** Floral traits of *Thelymitra* spp., *Calectasia* spp. and the 16 most abundant co-flowering species predicted to attract pollinators. Traits included are plant height, corolla height, corolla width, stamen length (in orchids, pseudoanther length), presence of exposed pollen, presence of vertical projection of stamens (in orchids, pseudoanther), corolla symmetry, corolla shape, inflorescence, flower orientation, and type of reward. Plants are categorised as 'orchid', 'putative model', 'other rewarding species' and species 'sharing colour' with the orchids. Corolla symmetry: zygomorphic (0) actinomorphic (1); corolla shape: rotate (r); papilionaceous (p); bilabiate (b); ligulate (l); flower orientation: upright (u); horizontal (h); anthers exposure: no (0) yes (1); vertical projection of stamens (in orchids pseudo anther), no (0) yes (1); inflorescence: umbel (u), raceme (r), spike (s), panicle (p), solitary (so).

| **Species** | **Categories** | **Plant height mean (mm)** | **Flower height (mm)** | **Flower wide (mm)** | **Stamen length (mm)** | **Exposed anthers** | **Vertical projection of stamens** | **Corolla symmetry** | **Corolla shape** | **Inflorescence** | **Flower orientation** | **Food reward** |
|---|---|---|---|---|---|---|---|---|---|---|---|---|
| *Acacia browniana* | other rewarding sp. | 1100.00 | 3.99 | 3.99 | 3.20 | 1 | 0 | 1 | r | p | h | pollen |
| *Acacia pulchella* | other rewarding sp. | 900.00 | 3.99 | 3.99 | 3.50 | 1 | 0 | 1 | r | p | h | pollen |
| *Actinodium cunninghamii* | other rewarding sp. | 600.00 | 25.32 | 25.32 | 1.00 | 1 | 0 | 1 | r | so | u | both |
| *Adenanthos obovatus* | other rewarding sp. | 900.00 | 28.08 | 7.76 | 1.50 | 1 | 0 | 0 | t | s | u | both |
| *Andersonia caerulea* | sharing colour | 500.00 | 6.36 | 6.77 | 0.00 | 0 | 1 | 1 | r | r | u | pollen |
| *Andersonia simplex* | sharing colour | 300.00 | 8.48 | 8.81 | 0.00 | 0 | 1 | 1 | r | s | u | both |
| *Calectasia gracilis* | putative model | 170.10 | 16.47 | 15.88 | 4.02 | 1 | 1 | 1 | r | r | u | pollen |
| *Calectasia demarzii* | putative model | 280.00 | 20.76 | 20.15 | 5.04 | 1 | 1 | 1 | r | r | u | pollen |
| *Conospermum caeruleum* | sharing colour | 650.00 | 1.92 | 1.84 | 0.80 | 0 | 0 | 0 | b | u | h | likely both |
| *Dampiera linearis* | sharing colour | 300.00 | 24.32 | 26.70 | 0.00 | 0 | 1 | 0 | r | r | u | likely both |
| *Daviesia hakeoides* | other rewarding sp. | 600.00 | 7.86 | 8.09 | 0.00 | 0 | 0 | 0 | p | p | h | both |
| *Hakea cucculata* | other rewarding sp. | 2500.00 | 26.39 | 26.37 | 4.50 | 1 | 0 | 0 | l | s | u | both |
| *Hakea lasiantha* | other rewarding sp. | 1800.00 | 5.60 | 9.78 | 1.40 | 1 | 0 | 0 | l | s | u | both |
| *Hibbertia psilocarpa* | other rewarding sp. | 600.00 | 11.94 | 16.94 | 1.50 | 1 | 0 | 1 | r | r | u | pollen |

| Species | Categories | Plant height mean (mm) | Flower height (mm) | Flower wide (mm) | Stamen length (mm) | Exposed anthers | Vertical projection of stamens | Corolla symmetry | Corolla shape | Inflorescence | Flower orientation | Food reward |
|---|---|---|---|---|---|---|---|---|---|---|---|---|
| *Isopogon formosus* | other rewarding sp. | 1100.00 | 60.00 | 60.00 | 4.00 | 1 | 0 | 1 | r | so | h | both |
| *Leucopogon distans* | other rewarding sp. | 700.00 | 8.48 | 8.81 | 0.00 | 0 | 0 | 1 | r | u | u | likely both |
| *Leucopogon glabellus* | other rewarding sp. | 600.00 | 4.53 | 4.53 | 0.00 | 0 | 0 | 1 | r | s | u | likely both |
| *Malaleuca suberosa* | sharing colour | 400.00 | 7.99 | 8.72 | 20.00 | 1 | 1 | 1 | r | s | u | both |
| *Petrophile squamata* | other rewarding sp. | 1650.00 | 15.98 | 15.92 | 4.00 | 1 | 0 | 0 | l | s | u | both |
| *Thelymitra pophyrostricta* | orchid | 230.4 | 38.379 | 38.481 | 5.26 | 1 | 1 | 1 | r | r | u | na |
| *Thelymitra speciosa* | orchid | 120.75 | 30.77 | 26.39 | 4.72 | 1 | 1 | 1 | r | r | u | na |

**Table S3.** Observations of floral visitors (AP=*Anthoglossa plumosa*; L=*Leioproctus* sp., TE=*Thricocolletes erythrinus*, LP=*Lasioglossum parasphecoides*, AM=*Apis mellifera*) on the two orchid species, *Thelymitra speciosa* and *T. porphyrosticta*, and the two tinsel lilies, *Calectasia gracilis* and *C. demarzii*. Number of individuals are given, with F indicating a female bee (carrying pollen loads on tibiae). In addition, behaviour when approaching the flower (DF=direct flight; ZF=zig-zag flight; PA=pointing antenna towards (false) anther), and when landing on the flower (B=bending body around (false) anthers; M=holding anthers with forelegs and attempting to manipulate anthers with mandibles, suggestive of buzz pollination; NF=nectar foraging attempt, pointing the head towards the stigmatic area) are indicated.

| | ***Thelymitra speciosa*** | | | | | ***Thelymitra porphyrosticta*** | | | | | ***Calectasia gracilis*** | | ***Calectasia demarzii*** | |
|---|---|---|---|---|---|---|---|---|---|---|---|---|---|---|
| **Number of insects/Behaviour** | ***AP*** | ***L*** | ***TE*** | **Beetle** | **Ant** | ***AP*** | ***L*** | ***LP*** | ***AM*** | **Ant** | **AP** | **L** | **AP** | **L** |
| Approaching the flower | 3 F | 1 F | 1 F | 2 | 1 | 2 F | 2 | 1 | 5 | 1 | 6 F | 4 | 5 F | 15 |
| Behaviour when approaching the flower | DF, PA | DF, PA | DF | ZF | ZF | DF, PA | DF, PA | ZF | ZF | ZF | DF, PA | DF, PA | DF, PA | DF, PA |
| Carrying orchid pollen on arrival | 0 | 0 | 0 | 0 | 0 | 0 | 0 | 0 | 0 | 0 | 4 | 3 F | 4 | 12 F |
| Landing on the flower | 2 F | 1 F | 1 | 3 | 1 | 2 | 2 F | 0. | 5 | 1 | 4 | 4 F | 5 | 15 F |
| Behaviour when landing on the flower | B, M | B, M | NF | NF | NF | B, M | B, M, NF | NF | NF | NF | B, M | B, M | B, M | B, M |
| Attempting false anther manipulation | 2 F | 1 F | 0 | 0 | 0 | 2 | 1 F | 0 | 0 | 0 | 4 | 4 F | 4 | 13 F |
| Removing pollen | 1 | 0 | 0 | 0 | 0 | 1 F | 0 | 1 | 0 | 0 | 4 F | 4 | 4 F | 13 |
| Depositing pollen | 0 | 0 | 0 | 0 | 0 | 0 F | 0 | 0 | 0 | 0 | 3 F | 2 | 4 F | 8 |
| Visiting another orchid flower | 1 | 0 | 0 | 1 | 0 | 1 F | 0 | 0 | 2 | 0 | 6 F | 4 | 5 F | 15 |

**Table S4.** List of insects caught on *Thelymitra* spp. and on co-occurring plants. All insects were sexed (F: female; M: male; W: worker) and identified to the species level (if possible) at the Western Australia Museum.

| Site | Coordinates | Date | WA Museum Voucher Code | Species | Family | where caught | Sex |
|---|---|---|---|---|---|---|---|
| Bremer Bay | -34.343641, 119.196235 | 14/08/2020 | WAM BBDA01 | *Trichocolletes erythrurus* | Colletidae | *Daviesia hakeoides* | F |
| Bremer Bay | -34.343641, 119.196235 | 14/08/2020 | WAM BBDA01 | *Trichocolletes erythrurus* | Colletidae | *Daviesia hakeoides* | F |
| Bremer Bay | -34.343641, 119.196236 | 14/08/2020 | WAM BBTS01 | *Trichocolletes erythrurus* | Colletidae | *Thelymitra speciosa* | M |
| Bremer Bay | -34.343641, 119.196237 | 14/08/2020 | WAM BBDA02 | *Trichocolletes erythrurus* | Colletidae | *Daviesia hakeoides* | M |
| Bremer Bay | -34.343641, 119.196238 | 14/08/2020 | WAM BBDA03 | *Neophyllotocus sp.* | Scarabaeidae | *Daviesia hakeoides* | *na* |
| Bremer Bay | -34.343641, 119.196239 | 14/08/2020 | WAM BBDA03 | *Trichocolletes erythrurus* | Colletidae | *Daviesia hakeoides* | F |
| Bremer Bay | -34.343641, 119.196240 | 14/08/2020 | WAM BBDA04 | *Lipotriches australica* | Halictidae | *Daviesia hakeoides* | F |
| Bremer Bay | -34.343641, 119.196241 | 15/08/2020 | WAM BBLC01 | *Leioproctus sp.* | Colletidae | *Leucopogon* sp. | F |
| Waychinicup National Park | -34.8, 118.3 | 8/06/2021 | WAM BBTS02 | *Lassioglossum* (*Parasphecodes*) sp. | Halictidae | *Thelymitra speciosa* | F |
| Bremer Bay | -34.343641, 119.196241 | 25/07/2021 | WAM BBTS03 | *Leioproctus (Leioproctus*) sp. | Colletidae | *Thelymitra speciosa* | F |
| Bremer Bay | -34.343641, 119.196241 | 25/07/2021 | WAM BBCN01 | *Chrysopasta elegans* | Tachinidae | *Calectasia narragara* | n.a |
| Gull rock National Park | -35.005765, 118.022299 | 18/09/2022 | WAM GRLE01 | Nemestrinidae | Diptera | *Leucopogon sp* | na |
| Gull rock National Park | -35.005765, 118.022300 | 18/09/2022 | WAM GRDA01 | *Trichocolletes erythrurus* | Colletidae | *Daviesia hakeoides* | M |
| Gull rock National Park | -35.005765, 118.022301 | 18/09/2022 | WAM GRLE02 | *Leioproctus* (*Leioproctus*) sp. | Colletidae | *Leucopogon sp* | F |
| Gull rock National Park | -35.005765, 118.022302 | 18/09/2022 | WAM GRDA02 | *Leioproctus* (*Leioproctus*) sp. | Colletidae | *Daviesia hakeoides* | F |
| Waychinicup National Park | -34.865122, 118.350503 | 18/09/2022 | WAM WYPE01 | *Lasioglossum* (*Chilalictus*) sp. | Halictidae | *Petrophile sp.* | F |
| Waychinicup National Park | -34.8, 118.3 | 18/09/2022 | WAM WYPE02 | *Leioproctus* (*Leioproctus*) sp. | Colletidae | *Isopogon formosus* | F |
| Waychinicup National Park | -34.8, 118.4 | 18/09/2022 | WAM WYPE03 | *Leioproctus* (*Leioproctus*) sp. | Colletidae | *Isopogon formosus* | F |
| Waychinicup National Park | -34.8, 118.5 | 18/09/2022 | WAM WYPE04 | *Leioproctus* (*Leioproctus*) sp. | Colletidae | *Isopogon formosus* | F |
| Boxwood Hill Ongerup Road | -34.8, 118.6 | 18/09/2022 | WAM BHPE01 | *Leioproctus* (*Leioproctus*) sp. | Colletidae | *Calectasia* sp. | F |
| Boxwood Hill Ongerup Road | -34.8, 118.7 | 19/09/2022 | WAM BHPE02 | *Leioproctus* (*Leioproctus*) sp. | Colletidae | *Calectasia* sp. | F |
| Gull Rock National Park | -35.005765, 118.022299 | 18/09/2022 | not vauchered | *Apis mellifera* | Apidae | *Daviesia hakeoides* | W |
| Gull Rock National Park | -35.005765, 118.022300 | 18/09/2022 | not vauchered | *Apis mellifera* | Apidae | *Andersonia caerulea* | W |
| Stirling Range National Park | -34.3329, 118.1214 | 18/08/2023 | WAM E114817 | *Anthoglossa plumosa* | Colletidae | *Calectasia* sp. | F |
| | | | **Pre-existing records** | | | | |
| Neerabup National Park | -31.6294, 115.7067 | 13-Aug-00 | WAM 27796 | *Anthoglossa* sp. M409/F471 | Colletidae | *Calectasia* sp. | M |
| Neerabup National Park | -31.6294, 115.7067 | 13-Aug-00 | WAM 27797 | *Anthoglossa* sp. M409/F471 | Colletidae | *Calectasia* sp. | M |
| Neerabup National Park | -31.6294, 115.7067 | 13-Aug-00 | WAM 27798 | *Anthoglossa* sp. M409/F471 | Colletidae | *Calectasia* sp. | M |
| Neerabup National Park | -31.6294, 115.7067 | 13-Aug-00 | WAM 27799 | *Anthoglossa* sp. M409/F471 | Colletidae | *Calectasia* sp. | F |
| Neerabup National Park | -31.6294, 115.7067 | 13-Aug-00 | WAM 27800 | *Anthoglossa* sp. M409/F471 | Colletidae | *Calectasia* sp. | F |
| Torndirrup National Park | -35.0952 S, 117.8664 | 25/05/1983 to 09/11/1983 | WAM 7718 | *Anthoglossa* sp. F114 | Colletidae | pitfall trap | F |
| Torndirrup National Park | -35.0952 S, 117.8664 | 25/05/1983 to 09/11/1983 | WAM 7719 | *Anthoglossa* sp. F114 | Colletidae | pitfall trap | F |
| Torndirrup National Park | -35.0952 S, 117.8664 | 25/05/1983 to 09/11/1983 | WAM 7720 | *Anthoglossa* sp. F114 | Colletidae | pitfall trap | F |
| Torndirrup National Park | -35.0952 S, 117.8664 | 25/05/1983 to 09/11/1983 | WAM 7721 | *Anthoglossa* sp. F114 | Colletidae | pitfall trap | F |

.

**Table S5.** List of the plots of *Thelymitra speciosa* and *T. porphyrosticta* populations surveyed in 2020, 2021 and 2022. All values are related to plants surveyed within 10 x 10 meter squares.

| **Date** | **Year** | **Species** | **Quadrat number** | **Population code** | **Site** | **Latitude, longitude** | **Mean pollen removal** | **Mean fruit production** | **N total flowers of *Calectasia*** | **N total flowers of *Daviesia hakeoides*** | **N total flowers of *Malalauca suberosa*** | **N total flowers of *Dampiera linearis*** | **N total flowers of *Andersonia*** | **N total flowers of *Conospermum caeruleum*** | **N total flowers of *Leucopogon glabellus*** | **N total flowers of *Leucopogon distans*** | **N total flowers of *Petrophile squamata*** | **N total flowers of *Hakea cucculata*** | **N total flowers of *Leptospermum laevigatum*** | **N total flowers with similar coloration** |
|---|---|---|---|---|---|---|---|---|---|---|---|---|---|---|---|---|---|---|---|---|
| 14/08/2020 | 2020 | *Thelymitra speciosa* | 1 | TZ1 | Tozer campsite | 34°20'27.7"S 119°11'42.8"E | 0.09 | 0.02 | 40 | 105 | 40 | 0 | 0 | 0 | 0 | 0 | 0 | 0 | 0 | 80 |
| 14/08/2020 | 2020 | *Thelymitra speciosa* | 2 | TZ2 | Tozer campsite | 34°20'36.4"S 119°11'44.7"E | 0.91 | 0.00 | 15 | 0 | 0 | 0 | 185 | 0 | 0 | 0 | 0 | 0 | 0 | 200 |
| 14/08/2021 | 2021 | *Thelymitra speciosa* | 3 | TZ1 | Tozer campsite | 34°20'36.4"S 119°11'44.7"E | 0.00 | 0.06 | 36 | 360 | 35 | 35 | 0 | 0 | 0 | 0 | 0 | 0 | 0 | 106 |
| 14/08/2021 | 2021 | *Thelymitra speciosa* | 4 | TZ2 | Tozer campsite | 34°20'34.6"S 119°11'42.4"E | 0.14 | 0.08 | 64 | 0 | 10 | 15 | 28 | 0 | 0 | 0 | 0 | 0 | 0 | 117 |
| 14/08/2021 | 2021 | *Thelymitra speciosa* | 5 | TZ3 | Tozer campsite | 34°20'27.7"S 119°11'42.8"E | 0.17 | 0.00 | 0 | 0 | 14 | 20 | 0 | 0 | 0 | 0 | 0 | 0 | 0 | 34 |
| 14/08/2021 | 2021 | *Thelymitra speciosa* | 6 | TZ4 | Tozer campsite | 34°20'24.8"S 119°11'35.7"E | 0.19 | 0.00 | 0 | 345 | 180 | 4 | 0 | 0 | 0 | 0 | 0 | 0 | 0 | 184 |
| 18/08/2021 | 2021 | *Thelymitra porphyrosticta* | 7 | WAY3 | Waychinicup NP | 34°52'08.5"S 118°21'11.9"E | 0.38 | 0.24 | 0 | 0 | 0 | 0 | 50 | 40 | 0 | 0 | 0 | 0 | 0 | 90 |
| 18/08/2021 | 2021 | *Thelymitra porphyrosticta* | 8 | WAY3 | Waychinicup NP | 34°52'01.1"S 118°21'06.8"E | 0.19 | 0.03 | 0 | 0 | 0 | 0 | 50 | 40 | 0 | 0 | 0 | 0 | 0 | 90 |
| 18/08/2021 | 2021 | *Thelymitra porphyrosticta* | 9 | WAY3 | Waychinicup NP | 34°51'56.0"S 118°21'02.0"E | 0.08 | 0.17 | 0 | 0 | 0 | 0 | 110 | 35 | 0 | 0 | 0 | 0 | 0 | 145 |
| 18/08/2021 | 2021 | *Thelymitra porphyrosticta* | 10 | WAY3 | Waychinicup NP | 34°51'50.8"S 118°20'58.8"E | 0.21 | 0.20 | 40 | 0 | 35 | 0 | 85 | 0 | 0 | 0 | 0 | 0 | 0 | 160 |
| 20/08/2021 | 2021 | *Thelymitra porphyrosticta* | 11 | GR4 | Gull rock NP | 35°00'22.2"S 118°01'23.0"E | 0.04 | 0.08 | 0 | 0 | 0 | 0 | 90 | 0 | 0 | 100 | 0 | 0 | 0 | 90 |
| 20/08/2021 | 2021 | *Thelymitra porphyrosticta* | 12 | GR4 | Gull rock NP | 35°00'22.6"S 118°01'22.9"E | 0.20 | 0.25 | 0 | 0 | 0 | 0 | 160 | 0 | 90 | 0 | 0 | 0 | 0 | 160 |
| 20/08/2021 | 2021 | *Thelymitra porphyrosticta* | 13 | GR4 | Gull rock NP | 35°00'21.6"S 118°01'20.4"E | 0.13 | 0.05 | 0 | 0 | 0 | 0 | 120 | 0 | 70 | 0 | 0 | 0 | 0 | 120 |
| 20/08/2021 | 2021 | *Thelymitra porphyrosticta* | 14 | GR4 | Gull rock NP | 35°00'23.1"S 118°01'21.3"E | 0.13 | 0.46 | 0 | 0 | 0 | 0 | 100 | 0 | 110 | 0 | 0 | 0 | 500 | 100 |
| 28/08/2022 | 2022 | *Thelymitra speciosa* | 15 | TZ1 | Tozer campsite | 34°20'36.4"S 119°11'44.7"E | 0.08 | 0.13 | 45 | 205 | 125 | 20 | 0 | 0 | 0 | 0 | 0 | 0 | 0 | 190 |
| 28/08/2022 | 2022 | *Thelymitra speciosa* | 16 | TZ3 | Tozer campsite | 34°20'34.6"S 119°11'42.4"E | 0.00 | 0.67 | 130 | 120 | 260 | 4 | 0 | 0 | 0 | 0 | 0 | 0 | 0 | 394 |
| 28/08/2022 | 2022 | *Thelymitra speciosa* | 17 | TZ4 | Tozer campsite | 34°20'27.7"S 119°11'42.8"E | 0.11 | 0.17 | 0 | 240 | 180 | 0 | 0 | 0 | 0 | 0 | 0 | 0 | 0 | 180 |
| 28/08/2022 | 2022 | *Thelymitra speciosa* | 18 | TZ2 | Tozer campsite | 34°20'24.8"S 119°11'35.7"E | 0.10 | 0.10 | 45 | 0 | 40 | 6 | 0 | 0 | 0 | 0 | 0 | 0 | 0 | 91 |
| 24/08/2022 | 2022 | *Thelymitra porphyrosticta* | 19 | WAY3 | Waychinicup NP | 34°52'08.5"S 118°21'11.9"E | 0.03 | 0.19 | 0 | 0 | 0 | 0 | 20 | 10 | 0 | 0 | 50 | 0 | 0 | 30 |
| 24/08/2022 | 2022 | *Thelymitra porphyrosticta* | 20 | WAY3 | Waychinicup NP | 34°52'01.1"S 118°21'06.8"E | 0.00 | 0.00 | 0 | 0 | 0 | 0 | 0 | 0 | 55 | 0 | 0 | 0 | 0 | 0 |
| 24/08/2022 | 2022 | *Thelymitra porphyrosticta* | 21 | WAY3 | Waychinicup NP | 34°51'56.0"S 118°21'02.0"E | 0.00 | 0.42 | 0 | 0 | 0 | 0 | 75 | 110 | 30 | 0 | 20 | 0 | 0 | 185 |
| 24/08/2022 | 2022 | *Thelymitra porphyrosticta* | 22 | WAY3 | Waychinicup NP | 34°51'50.8"S 118°20'58.8"E | 0.00 | 0.00 | 0 | 0 | 0 | 0 | 0 | 0 | 15 | 30 | 0 | 0 | 0 | 0 |
| 26/08/2022 | 2022 | *Thelymitra porphyrosticta* | 23 | GR5 | Gull rock NP | 35°00'14.9"S 118°01'05.2"E | 0.10 | 0.00 | 0 | 0 | 0 | 0 | 90 | 0 | 60 | 0 | 0 | 0 | 0 | 90 |
| 26/08/2022 | 2022 | *Thelymitra porphyrosticta* | 24 | GR6 | Gull rock NP | 35°00'27.3"S 118°01'44.3"E | 0.03 | 0.04 | 0 | 0 | 0 | 0 | 120 | 120 | 200 | 0 | 0 | 0 | 1000 | 240 |
| 26/08/2022 | 2022 | *Thelymitra porphyrosticta* | 25 | GR7 | Gull rock NP | 35°00'26.1"S 118°02'06.3"E | 0.20 | 0.16 | 0 | 0 | 0 | 0 | 120 | 0 | 200 | 0 | 0 | 0 | 1000 | 120 |
| 26/08/2022 | 2022 | *Thelymitra porphyrosticta* | 26 | GR4 | Gull rock NP | 35°00'37.1"S 118°02'11.9"E | 0.11 | 0.21 | 0 | 0 | 0 | 0 | 140 | 0 | 180 | 0 | 0 | 0 | 0 | 140 |

**Table S6.** Compounds with compound class present in the orchids *Thelymitra speciosa* and *T. porphyrosticta*, and in co-flowering plants. Compound retention indices (RI) calculated for the compounds and reported in the Literature for DB-5 GC column. *Confirmed with commercial standard ^DB-5MS instead of DB-5.

| | | | | | Relative abundance (%) | | | | | | | | | |
|---|---|---|---|---|---|---|---|---|---|---|---|---|---|---|
| **Compound** | **Class** | **RI (DB-5)** | **RI (Literature)** | **Reference** | ***Thelymitra speciosa*** | ***Thelymitra porphyrostricta*** | ***Calectasia gracilis*** | ***Calectasia demarzii*** | ***Malaleuca suberosa*** | ***Andersonia caerulea*** | ***Daviesia hakeoides*** | ***Leucopogon glabellus*** | ***Leptospermum laevigatum*** | ***Hibbertia psilocarpa*** |
| Limonene * | Monoterpene | 1039 | 1039 | Choi, 2003 | 23.40 | 6.89 | 21.26 | - | 8.66 | 3.96 | - | - | 14.47 | - |
| β-Pinene * | Monoterpene | 989 | 989 | Buchin *et al.*, 2002 | 3.16 | - | 2.67 | - | - | 2.75 | - | - | 38.79 | - |
| Acetic Acid, hexyl ester | Carboxylic acid ester | 1010 | 1011 | Beaulieu and Grimm, 2001; Mayr *et al.*, 2003 | 3.48 | - | 2.33 | - | - | - | - | - | - | - |
| p-Cymene * | Monoterpene | 1034 | 1034; 1033 | Buchin *et al.*, 2002; Hognadottir, 2003 | 4.95 | - | 4.32 | - | - | - | - | - | 3.12 | - |
| Eucalyptol * | Monoterpenoid | 1044 | 1044; 1048 | Buchin *et al.*, 2002; Baranauskiene *et al.*, 2003 | 1.75 | - | 1.08 | - | 64.62 | - | - | - | 31.26 | - |
| γ-Terpinene * | Monoterpenoid | 1067 | 1067 | Loayza *et al.*, 1995; Merle *et al.*, 2004 | 3.71 | - | 3.19 | - | - | - | - | - | - | - |
| Terpinolene | Monoterpenoid | 1096 | 1098; 1096 | Merle *et al.*, 2004; Högnadóttir and Rouseff, 2003 | 3.71 | - | 1.53 | - | - | - | - | - | - | - |
| Hexanoic acid, butyl ester | Saturated fatty acid ester | 1189 | 1189; 1189^ | Fan and Qian, 2006; Maia *et al.*, 2002 | 2.24 | - | 1.34 | - | - | - | - | - | - | - |
| α-Pinene | Monoterpene | 941 | 940 | Lucero *et al.*, 2006 | - | - | - | - | 2.28 | 5.55 | - | - | - | - |
| Myrcene | Monoterpene | 990 | 991 | Adams, 1995; Choi, 2003 | - | - | - | - | 10.41 | - | - | - | - | - |
| Camphene | Monoterpene | 941 | 948 | Da Silva *et al.*, 1999 | - | - | - | - | - | 16.62 | - | - | - | - |
| 1-Heptanol | Fatty alcohol | 968 | 967; 969 | Rostad and Pereira, 1986; Beaulieu and Grimm, 2001 | - | - | | - | - | - | 5.77 | 5.37 | - | - |
| Decane | Hydrocarbon | 999 | 999 | Adams, 1995 | - | - | - | - | - | - | 2.24 | 21.81 | 4.58 | 4.89 |
| (E)-p-mentha-2,8-dien-1-ol | Monoterpenoid | 1122 | 1122 | https://www.flavornet.org/f_kovats.html | - | - | - | - | - | - | - | - | 2.85 | - |
| Verbenone | Monoterpene | 1222 | 1218 | Hognadottir, 2003 | - | - | - | - | - | - | - | - | 2.76 | - |
| (E)-Caryophyllene | Sequiterpene | 1435 | 1433 | Mosayebi *et al.*, 2008 | - | - | - | - | - | - | - | - | 2.48 | - |
| Unknown $C_{15}H_{30}$ | - | 1487 | - | - | 4.57 | 23.56 | 3.7 | - | - | - | - | - | - | - |
| Unknown $C_{11}H_{18}O_2$ | - | 1491 | - | - | 20.00 | 31.57 | 22.49 | 8.47 | - | - | - | - | - | - |
